# Manipulating the 3D organization of the largest synthetic yeast chromosome

**DOI:** 10.1101/2022.04.09.487066

**Authors:** Weimin Zhang, Luciana Lazar-Stefanita, Hitoyoshi Yamashita, Michael J. Shen, Leslie A. Mitchell, Hikaru Kurasawa, Evgenii Lobzaev, Viola Fanfani, Max A. B. Haase, Xiaoji Sun, Qingwen Jiang, Gregory W. Goldberg, David M. Ichikawa, Stephanie L. Lauer, Laura H. McCulloch, Nicole Easo, S. Jiaming Lin, Brendan R. Camellato, Yinan Zhu, Jitong Cai, Zhuwei Xu, Yu Zhao, Maya Sacasa, Ryan Accardo, Ju Young Ahn, Surekha Annadanam, Leighanne A. Brammer Basta, Nicholas R. Bello, Lousanna Cai, Stephanie Cerritos, MacIntosh Cornwell, Anthony D’Amato, Maria Hacker, Kenneth Hersey, Emma Kennedy, Ardeshir Kianercy, Dohee Kim, Hong Seo Lim, Griffin McCutcheon, Kimiko McGirr, Nora Meaney, Lauren Meyer, Ally Moyer, Maisa Nimer, Carla Sabbatini, Lucas S. Shores, Cassandra Silvestrone, Arden Snee, Antonio Spina, Anthony Staiti, Matt Stuver, Elli Tian, Danielle Whearty, Calvin Zhao, Tony Zheng, Vivian Zhou, Lisa Scheifele, Karen Zeller, Marcus B. Noyes, Joel S. Bader, Samuel Deutsch, Giovanni Stracquadanio, Yasunori Aizawa, Junbiao Dai, Jef D. Boeke

**Affiliations:** Institute for Systems Genetics and Department of Biochemistry and Molecular Pharmacology, NYU Langone Health, New York, NY, USA; Department of Biomedical Engineering, NYU Tandon School of Engineering, Brooklyn, NY, USA; School of Life Science and Technology, Tokyo Institute of Technology, Yokohama, Japan; School of Biological Sciences, The University of Edinburgh, Edinburgh, UK; CAS Key Laboratory of Quantitative Engineering Biology, Guangdong Provincial Key Laboratory of Synthetic Genomics and Shenzhen Key Laboratory of Synthetic Genomics, Shenzhen Institute of Synthetic Biology, Shenzhen Institute of Advanced Technology, Chinese Academy of Sciences, Shenzhen, China; Department of Biomedical Engineering, Whiting School of Engineering, Johns Hopkins University, Baltimore, MD, USA; Department of Biology, Loyola University Maryland, 4501 N Charles St, Baltimore, MD, USA; Department of Biology, Krieger School of Arts and Sciences, Johns Hopkins University, Baltimore, MD, USA; Brady Urological Institute, Johns Hopkins University School of Medicine, Baltimore, MD, USA; Department of Chemical and Biomolecular Engineering, Whiting School of Engineering, Johns Hopkins University, Baltimore, MD, USA; DOE Joint Genome Institute, Lawrence Berkeley National Laboratory, Berkeley, CA, USA; Kanagawa Institute of Industrial Science and Technology (KISTEC), Ebina, Kanagawa, 243-0435 Japan; Shenzhen Branch, Guangdong Laboratory for Lingnan Modern Agriculture, Key Laboratory of Synthetic Biology, Ministry of Agriculture and Rural Affairs, Agricultural Genomics Institute at Shenzhen, Chinese Academy of Agricultural Sciences, Shenzhen, China

**Keywords:** *Saccharomyces cerevisiae*, *synIV*, megachunk assembly, chromosome 3D structure manipulation, inside-out chromosome, chromosome tethering, transcriptomics

## Abstract

Whether synthetic genomes can power life has attracted broad interest in the synthetic biology field, especially when the synthetic genomes are extensively modified with thousands of designer features. Here we report *de novo* synthesis of the largest eukaryotic chromosome thus far, *synIV*, a 1,454,621-bp *Saccharomyces cerevisiae* chromosome resulting from extensive genome streamlining and modification. During the construction of *synIV*, we developed megachunk assembly combined with a hierarchical integration strategy, which significantly increased the accuracy and flexibility of synthetic chromosome construction and facilitated chromosome debugging. In addition to the drastic sequence changes made to *synIV* by rewriting it, we further manipulated the three-dimensional structure of *synIV* in the yeast nucleus to explore spatial gene regulation within the nuclear space. Surprisingly, we found few gene expression changes, suggesting that positioning inside the yeast nucleoplasm plays a minor role in gene regulation. Lastly, we tethered *synIV* to the inner nuclear membrane via its hundreds of *loxPsym* sites and observed transcriptional repression of the entire chromosome, demonstrating chromosome-wide transcription manipulation without changing the DNA sequences. Our manipulation of the spatial structure of the largest synthetic yeast chromosome shed light on higher-order architectural design of the synthetic genomes.

**Graphical abstract:** 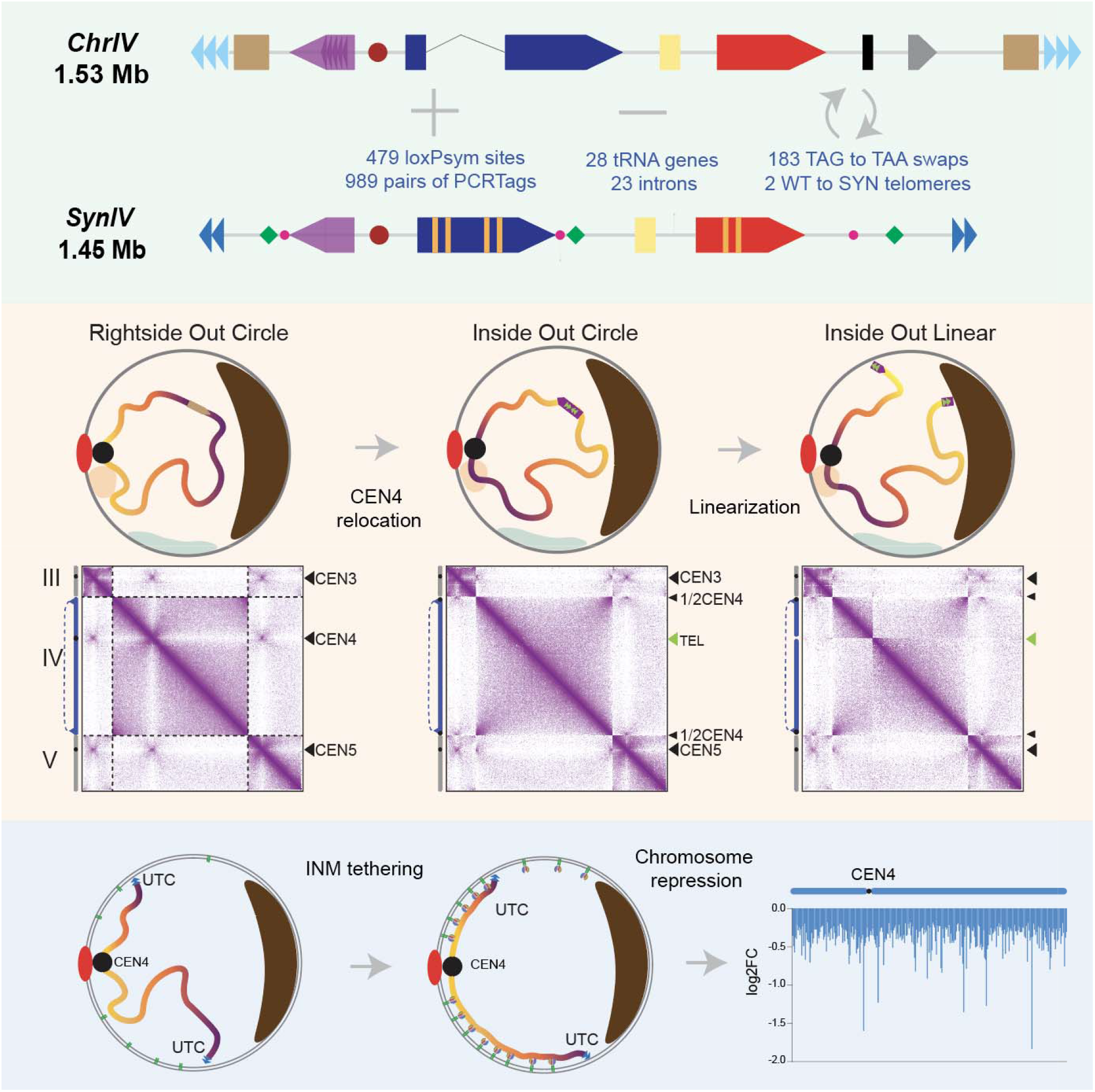

*Highlights:*

- *De novo* synthesis of the largest eukaryotic chromosome, *synIV*
- *SynIV* shows similar 3D structure to wild-type IV, despite thousands of changes made to it
- “Inside-out” repositioning of *synIV* in nucleus shows minor transcriptional changes
- Multipoint tethering *synIV* to inner nuclear membrane represses transcription of whole chromosome

## Introduction

Reprogramming an entire native genome facilitates the understanding of many fundamental questions regarding genome content essentiality, genome regulation and genome evolution^1,2^ while also increasing its versatility by incorporating new design features^3–5^. Although our ability to synthesize genomes has increased dramatically in the past two decades^6^, it is still challenging to accurately and efficiently build megabase-sized DNA segments in a bottom-up fashion, hindering studies aiming to probe the functional basis of genomes from a synthetic biology perspective. *Saccharomyces cerevisia*e was the first eukaryotic organism to have its genome sequenced, which profoundly facilitated genetic studies of yeast. Further, the Synthetic Yeast Genome Project (Sc2.0) represents the first and largest eukaryotic synthetic genome to be built. The project aims to drastically alter the yeast genome and assesses whether the native yeast genome can be reprogrammed by removing retrotransposons and other repetitive sequences in order to increase genome stability, whether splicing systems can be eliminated in an intron-less yeast genome and whether synthetic yeast can gain new properties when the entire gene content is “shuffled” randomly. Synthesis of the individual synthetic chromosomes have answered many fundamental questions^7–13^, yet much more remains open as Sc2.0 is driven to completion.

In the process of building Sc2.0 genome, the focus thus far has been on sequences of the chromosomes, and less so on how synthetic chromosomes are spatially organized in a living cell. Numerous studies have shown that genome misfolding and dysregulation are strongly associated with human disease^14–16^, suggesting fundamentally important roles of large scale chromatin 3D organization. It is crucial to have a thorough understanding of how synthetic chromosomes are organized in the nucleus so that we have a better control of chromosome-wide gene regulation. In yeast, chromosomes are organized in a relatively conserved structure called the Rabl orientation^17^–the spindle pole body (SPB) resides on the opposite side of the nucleus relative to the nucleolus and tethers the 16 centromeres throughout the cell cycle. The 32 telomeres are clustered to form three to eight foci anchored on the inner nuclear envelope^18,19^. Such nuclear organization forms different sub-nuclear compartments that influence genome-wide gene expression. For example, the peripheral domain of the nuclear pore complex corresponds to an “active expression” compartment, whereas telomere clusters on the inner nuclear membrane correspond to a repressive compartment. The nucleolus is paradoxical due to its nuclear apposition, where silencing is seen of RNAP2 reporter genes, but ribosomal DNA itself is actively transcribed by RNAP1 and RNAP3. However, to our knowledge, current findings are limited to compartments close to the nuclear envelope. It is still unclear whether the yeast *nucleoplasm* contains multiple, elaborately structured sub-nuclear compartments.

Here, following the completion of previous synthetic chromosomes (*synII*, *synIII*, *synV*, *synVI*, *synIXR*, *synX* and *synXII*), we describe the construction and characterization of a 1,454,641 bp yeast chromosome, *synIV*, the largest synthetic eukaryotic chromosome reported. Despite the incorporation of thousands of designer features, *synIV* is still able to provide near wild-type fitness to its host strain under various growth conditions. *SynIV* shows remarkably similar 3D structure to wild-type IV, and thanks to its lack of repetitive DNA, smoother intra-chromosomal contact maps are produced by chromosome conformation capture assay. Lastly, we redefined the 3D structure of *synIV* in the nucleus by “intramolecular centromere relocation.” The resulting *synIV* is flipped in arm orientation relative to the other 15 wild-type chromosomes, giving rise to the reorientation and repositioning of all 796 genes carried on *synIV* in the nucleus, and forcing centromere proximal sequences to telomere proximal and vice versa – producing an “inside out” chromosome. By manipulating the 3D organization of *synIV*, we observed surprisingly minor gene expression changes relative to the original *synIV*, indicating that *synIV* organization is highly plastic and suggesting that there are no functional subnuclear compartments that influence gene expression in the interior space of the yeast nucleoplasm. This finding leaves the nuclear envelope compartment as the only major repression site. By multipoint tethering *synIV,* along with other synthetic chromosomes, to the inner nuclear membrane, we achieved whole synthetic chromosome repression without altering the DNA sequences. Our work pushes the upper size boundary of synthetic eukaryotic chromosome construction and provides the first synthetic chromosomes pervasively engineered to adopt altered 3D conformations by design.

## Results

### Hierarchical assembly of *synIV*

Chromosome IV of *Saccharomyces cerevisia*e encodes the highest number of genes among the 16 chromosomes. Following the design pipeline of the Sc2.0 project^5^, the synthetic chromosome IV (*synIV*) becomes the largest eukaryotic chromosome to be built thus far. A total of 479 loxPsym (symmetrical loxP, which allows inversions and deletions in conjunction with any other loxPsym site with equal probability) sites were inserted downstream of non-essential genes and at the positions of deleted tRNA genes, 23 introns and 28 tRNA genes were removed, 183 TAG stop codons were swapped to TAA and 989 pairs of PCRTags were generated by synonymous recoding to serve as synthetic “watermarks” (**Figure 1A, Figure S1**)^7–13^. To minimize the possibility of genome instability, subtelomeric repeats and retrotransposons were removed, and seven open reading frames (ORFs) containing tandemly repeated regions^20^ were synonymously recoded (repeat smashed ORFs, **Table S1**).

**Fig. 1.**
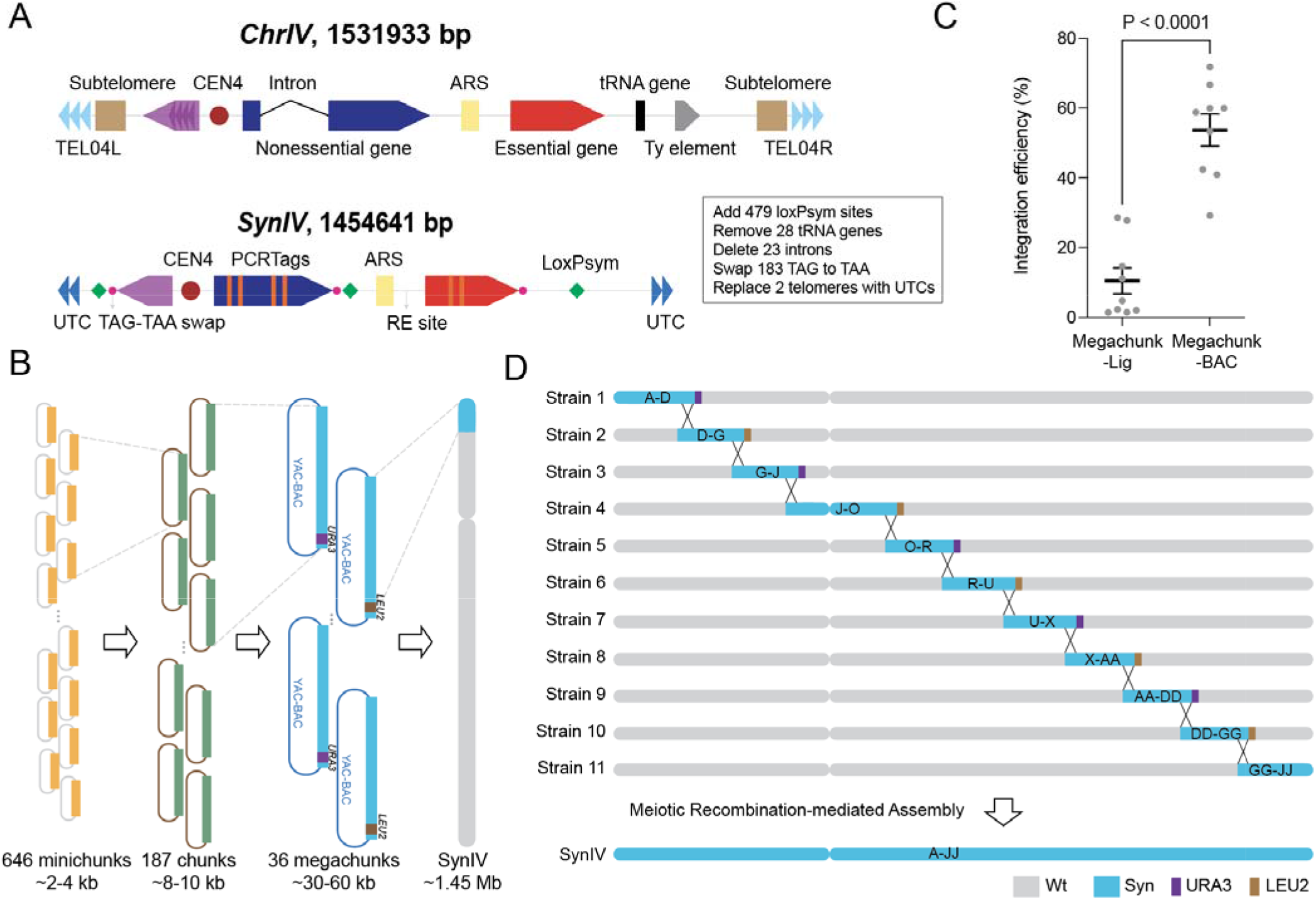
Design and assembly of *synIV*. (**A**) Schematic illustration of the individual types of design features of *synIV* compared to its native *chrIV* counterpart, including subtelomeric deletion; REPEATSMASHER recoding; TAG to TAA stop codon swaps (small red dots); PCRTag recoding (orange vertical bars); intron deletion; tRNA gene and adjacent Ty element deletion combined with loxPsym site insertion (green diamonds); *CEN*, centromere (large red dot); RE, restriction enzyme site; UTC, universal telomere cap (dark blue triangles); ARS, autonomously replicating sequence (yellow rectangle); native *TEL*, telomere (light blue triangles). Genes are categorized by different colors for nonessential gene (dark blue), repeat containing gene (purple), and essential gene (red). For full chromosome design, see Figure S1. (**B**) *SynIV* hierarchical assembly workflow. Yellow bars are ∼2-4 kb minichunk DNAs, green bars are ∼8-10 kb chunk DNAs, blue bars are ∼30-60 kb megachunk DNAs. YAC-BAC, yeast artificial chromosome-bacterial artificial chromosome shuttle vector. (**C**) Integration efficiency comparison between conventional ligation method and the new megachunk assembly method (n=9 for each method). Bars are mean±SEM. Integration efficiency was computed by number of successfully integrated clones (verified by presence of synthetic PCRTags and absence of wild-type PCRTags) divided by total number of clones screened. (**D**) Full length *synIV* was split into 11 intermediate strains for parallelization. Semi-*synIV* segments from different intermediate strains were consolidated via Meiotic Recombination-mediated Assembly (MRA). A *URA3* marker was originally built into megachunk JJ, and subsequently removed after obtaining the megachunk GG-JJ strain.

The 1,454,641 bp *in silico* designed *synIV* was segmented into 36 megachunks of approximately 60 kb in length (named megachunks A through JJ), which were further subdivided into 646 minichunks of about 3 kb each. Minichunks were assembled from oligonucleotides by DNA synthesis vendors. Adjacent minichunks share 100 bp of identity, allowing assembly of three to four minichunks into 8-10 kb DNA chunks (**Figure S2A**). Each chunk was flanked by two rare cutting restriction enzyme (RE) sites previously designed into ORFs by synonymous recoding (**Table S2**). Yeast homologous recombination or Gibson assembly were used to assemble the chunks (**Figure 1B**)^7,21^. Initially, five or six chunks were ligated *in vitro* to form a megachunk, and used directly for SwAP-In (<u>Sw</u>itching <u>A</u>uxotrophies <u>P</u>rogressively for <u>In</u>tegration), a methodology used to “overwrite” each wild-type chromosome segment with its synthetic counterpart^5^. We noted that the SwAP-In success rate varied considerably among different megachunk integrations due to inconsistent ligation efficiency/yield of the full length megachunk products, and furthermore, the restriction fragment end ligation accuracy was never validated prior to integration, leading to a higher-than-average mutation rate within the RE site junctions.

In order to standardize the synthetic DNA used for SwAP-In, we attempted to assemble megachunks as cloned building blocks for this chromosome (**Figure S2B, Table S3**). The megachunk assembly approach produced a significantly higher integration efficiency when compared to the previous method (**Figure 1C**) and, since all megachunk clones were sequenced prior to use, this improved methodology eliminated the problem of errors arising from RE end misligation. To accelerate the construction of *synIV*, we performed SwAP-In using 11 entry strains in parallel, among which any two adjacent entry strains have the opposite mating type and different auxotrophic markers. Ten of the 11 semi-synthetic strains have only four megachunks, while the megachunk J-O strain contains six, and each pair of adjacent strains shares a single overlapping megachunk. This design strategy facilitates Meiotic Recombination-mediated Assembly (MRA) in the next assembly phase (**Figure 1D**)^13^. After completing the integration of 11 semi-synthetic strains, we performed fitness characterization under various growth conditions (**Figures S3A-B**). We found that strains with G-J megachunks had a severe growth defect in rich medium (YPD) at high temperature; those with J-O megachunks grew poorly under all conditions we tested; while the strain with U-X megachunks showed low fitness in liquid cultures (**Figure S3B**). The unhealthy strains were subjected to debugging (see section below “Debugging of *synIV*”) until fitness met a near wild-type criterion. Karyotyping analysis of all semi-*synIV* chromosomes revealed no large-scale genomic rearrangement was created during the construction process (**Figure S3C**). These data suggest that megachunk cloning combined with hierarchical assembly strategy significantly increased the accuracy and flexibility of *synIV* construction.

### *SynIV* debugging

Characterization of semi-*synIV* strains revealed that strains containing megachunks G-J, J-O and U-X were defective in growth under at least one tested condition (**Figures S3A-B**). As the megachunks were incorporated in consecutive order, we characterized all strains generated at each SwAP-In step, searching for the defect(s) caused by megachunk insertion in each unhealthy strain. Eventually we narrowed down the bugs to megachunks J, N, O and V and classified them according to three types of alterations: repeat-smashed ORFs, PCRTag recoding, and unexpected mutations (**Figure 2A**).

**Fig. 2.**
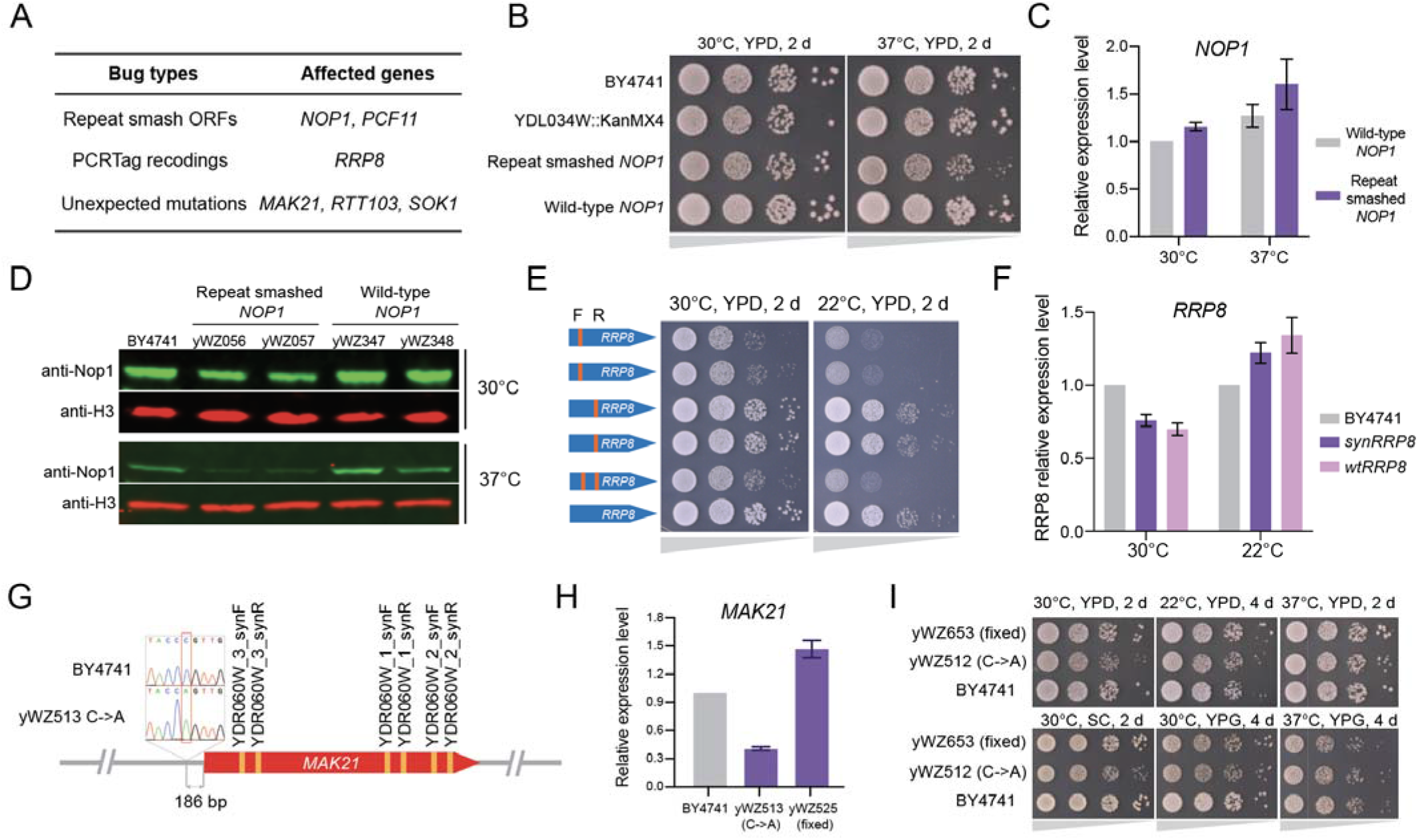
*SynIV* debugging. (**A**) Summary of bugs and genes in *synIV*. (**B**) Serial dilution spot assay of strains with different versions of *NOP1*. BY4741 and *YDL034W* knock-out (entry strain of megachunk J) strains served as controls. (**C**) RT-qPCR analysis of wild-type and repeat smashed *NOP1* mRNA expression at 30°C and 37°C. Bars represent mean ± SD of three technical replicates. (**D**) Immunoblotting analysis of Nop1p; two independent clones were used for each strain. (**E**) Serial dilution spot assay showing the fitness of strains with *RRP8* with only forward synthetic PCRTag, only reverse synthetic PCRTag or both forward and reverse synthetic PCRTags. F, forward synthetic PCRTag; R, reverse synthetic PCRTag. (**F**) Relative expression of synthetic and wild-type *RRP8* at 30°C and 22°C on YPD. Bars represent mean ± SD of three technical replicates. (**G**) Illustration of synthetic *MAK21* in *synIV*, a C to A mutation is highlighted in the promoter region of *MAK21.* Synthetic PCRTags are indicated as orange. (**H**) Relative expression of *MAK21* with or without the intergenic C->A mutation. Bars represent the mean ± SD of three technical replicates. (I) Serial dilution spot assay showing the growth fitness of strains with or without the intergenic C to A mutation.

Two *synIV* bugs resulted from the use of a piece of code, REPEATSMASHER^22^, designed to recode a set of a few dozen protein coding genes containing tandem repeat sequences that create challenges for assembly (**Table S1**). This version of REPEATSMASHER recoded the entire ORF rather than only the repetitive segment, resulting in a pervasively recoded ORF. For example, the “repeat-smashed” essential gene, *NOP1*, was found to cause a severe growth defect, especially when grown at 37°C on YPD (**Figure 2B**). This growth defect was fully reversed when repeat-smashed *NOP1* was replaced with wild-type *NOP1* (**Figure 2B**). We examined *NOP1* mRNA levels in repeat-smashed and wild-type strains and found that mRNA levels were not significantly different at both 30°C and 37°C YPD conditions (**Figure 2C**). However, the protein level of repeat-smashed Nop1p was substantially decreased at 37°C, and slightly decreased at 30°C (**Figure 2D**). Although the two *NOP1* genes encode identical amino acid sequences, they only share 74% identity at the DNA sequence level (**Figure S4A**). Importantly, repeat-smashed *NOP1* contains more non-optimal (based on codon usage frequency) codons for *S. cerevisiae* (**Figure S4B**), which may interfere with mRNA stability, translation initiation/elongation, and/or protein folding^23–25^. Thus, we speculate that the fitness defect is probably caused by altered translation of pervasively recoded *NOP1*. Similarly, the incorporation of *PCF11*, a repeat-smashed essential gene in megachunk V, caused growth defects under multiple conditions, which were rescued by reverting the repeat-smashed *PCF11* to the wild type (**Figure S4C**).

A PCRTag bug was discovered when megachunk O was integrated. The growth defect phenotype was especially notable when the strain was grown at low temperature (22°C) (**Figure S5A**). To narrow in on the bug, we integrated each chunk of megachunk O individually to a wild-type strain. We found that introducing chunk O4 alone to was sufficient to produce a similar fitness defect to that seen following whole megachunk O integration (**Figure S5B**). Further experiments mapped the bug to the sequence of the *RRP8* gene. The only unique feature of synthetic *RRP8* is a pair of PCRTags, and we eventually pinpointed the causative bug to the forward synthetic PCRTag, a 28-bp sequence in *RRP8* (**Figure 2E**). The incorporation of this 28-bp sequence did not compromise mRNA expression of *RRP8* (**Figure 2F**). mRNA secondary structure prediction indicates the PCRTag recoding produces a strong stem loop structure that may interfere with the translation of Rrp8p (**Figure S5C**), similar to the situation described for the *PRE4* gene on *synVI*^9^.

Some bugs were caused by unexpected (i.e. non-designed) mutations in the synthetic DNA (**Figure 2A**). In the case of the *MAK21* gene, we initially discovered that a segment containing *MAK21* was duplicated in megachunk J, and the duplication gave rise to better growth of megachunk J-N strain (**Figures S6A-B**). However, restoration of the *MAK21* coding sequence was unable to rescue the growth defect (**Figure S6C**). Instead, we discovered a single unplanned base-pair substitution, lying 186 bp upstream of *MAK21* in its promoter region (**Figure 2G**). This mutation resulted in a >50% reduction of *MAK21* mRNA expression (**Figures 2H-I**). Searching for transcription factor (TF) binding motif consensus sequences in *MAK21*’s promoter using the Yeastract+ tool^26^, we failed to identify any known TFs that bind to this −186 bp motif, suggesting that a novel functional element is disrupted by this point mutation. After correcting this bug and the aforementioned bugs, the 11 healthy semi-*synIV* strains were consolidated via multiple rounds of meiotic recombination assembly (MRA) (**Figure 3A**). The MRA combined synthetic chromosome segments efficiently until we approached the last step. We found that an *E. coli* IS1 transposon had inserted into *RTT103* in megachunk Y (**Figure S7A**), which did not cause any growth defect in strain yWZ462 (containing megachunk A-M and X-JJ) (**Figure S7B**), but led to severe growth defects on SC medium in strains containing almost all megachunks of *synIV* (**Figure S7C**, top3 rows). We fixed this “combinatorial” bug by removing the IS1 sequence from the *RTT103* gene, producing a healthy *synIV* strain (**Figure S7D**).

**Fig. 3.**
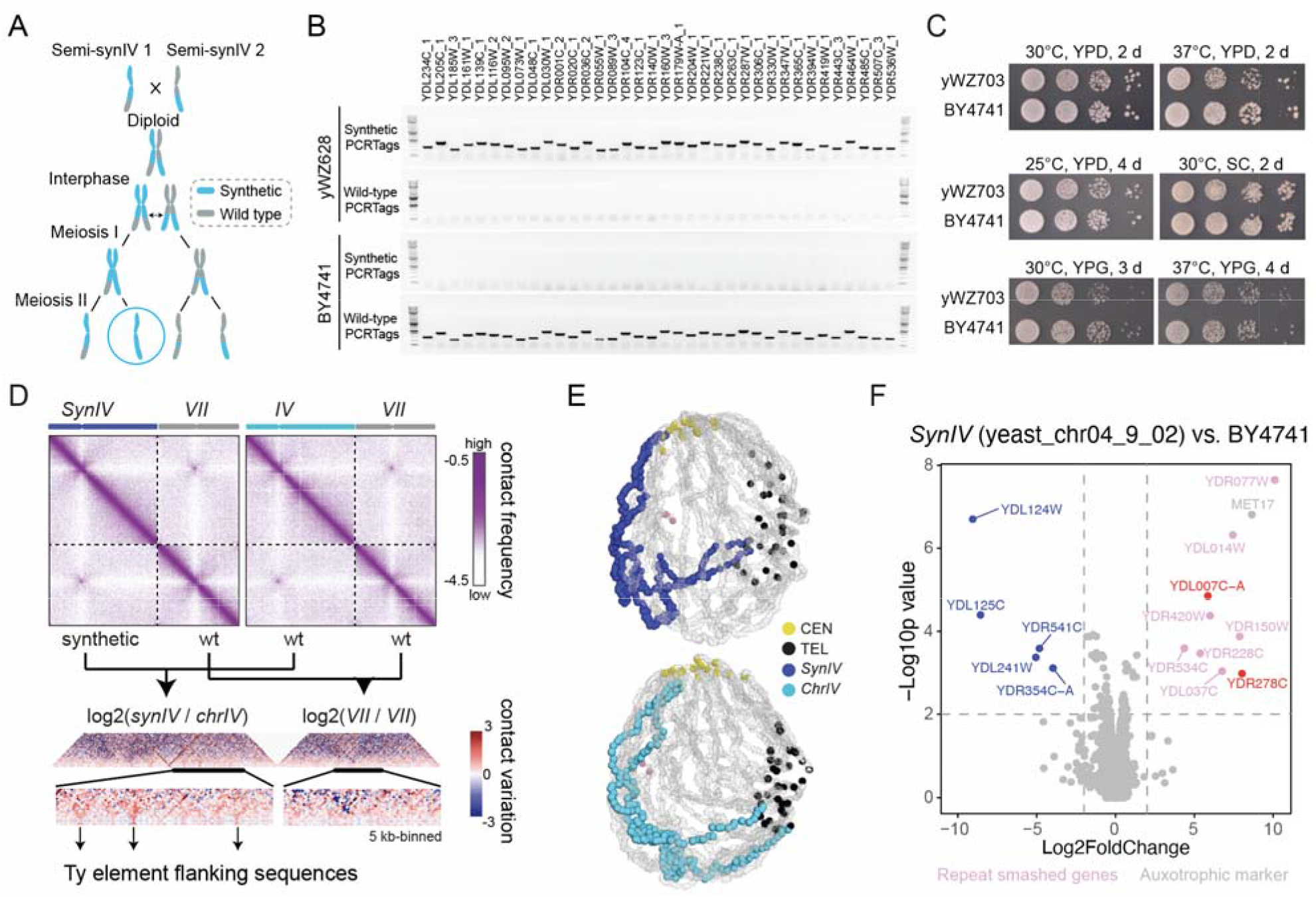
*SynIV* characterization. (**A**) Schematic illustration of Meiotic Recombination-mediated Assembly (MRA). (**B**) PCRTagging analysis of *synIV* (yWZ628) and wild-type (BY4741) strains. One PCRTag per megachunk was used for representation. Amplicons were loaded in 1.5% agarose gel for electrophoresis. NEB 1kb plus DNA ladder was used. (**C**) *SynIV* and BY4741 fitness comparison by ten-fold serial dilution spot assays under six growth conditions. YPD, yeast extract, peptone, dextrose. YPG, yeast extract, glycerol. SC, synthetic complete. (**D**) Hi-C contact maps of yeast with synthetic or wild-type chromosome IV. Chromosome VII is shown as an internal control. Chromosomes are annotated atop the maps in blue, cyan, and gray, respectively. Left and right maps (5 kb-binned) were generated by aligning Hi-C reads on a reference sequence containing the designed *synIV* chromosome. Violet to white color scale reflects high to low contact frequency (log10). Contact variations in *synIV* vs. wild-type IV are shown in the bottom panels where blue to red color scale reflects contact enrichment in *synIV* (log2-ratio maps 5 kb bin). (**E**) 3D average representations of the Hi-C contact maps in (D), color code highlights *synIV* and wild-type IV, in addition to centromeres, telomeres and rDNA. (**F**) Volcano plot of *synIV* (yeast_chr04_9_02) vs. BY4741. Genes are colored red for upregulated and blue down regulated in *synIV*, when compared to BY4741. Auxotrophic gene *MET17* was present in *synIV* but not in BY4741, whereas *YDL124W* and *YDL125C* were deleted in this yeast_chr04_9_04 *synIV* strain, but were restored in subsequent strain versions. Subtelomeric *YDL241W* was removed by design in *synIV*. Gray colored genes indicate genetic markers, pink genes that were repeat smashed, and light blue were 2-micron plasmid genes. Fold change cutoff is 4, p-value cutoff is 0.01.

### Sequence and structure of *synIV*

Meiotic recombination mediated assembly created a variety of intermediate *synIVs* that contained interspersed wild-type patches. Eventually, those intermediate *synIVs* were intercrossed to produce a full length *synIV* draft strain, named yeast_chr04_9_01. A PCR assay using selected synthetic and wild-type PCRTags showed that the wild-type chromosome IV was completely replaced by *synIV* (**Figure 3B**). Whole genome sequencing of yeast_chr04_9_01 revealed two structural variants: *ENA5-ENA2-ENA1* duplication and *HXT7-HXT6* duplication (**Figure S8A**). The *ENA* genes are greater than 90% identical to each other and are arranged in tandem in the genome, providing a favorable configuration for duplication in yeast. We successfully removed the duplicated *ENA* genes by first inserting a *URA3* gene near the duplicated region, and then replacing the duplicated region as well as the *URA3* gene with the correct DNA chunk. The *HXT7-HXT6* duplication is a well-known genome rearrangement event previously seen during evolution experiments under glucose limitation^27^, suggesting that the *synIV* strain may have experienced low glucose stress during construction. Besides these structural variants, 12 TAG stop codons were incorporated erroneously due to bookkeeping errors. We employed SpCas9-NG, a SpCas9 variant that recognizes the NG protospacer adjacent motif into yeast^28^ to precisely swap TAG stop codons to TAA. The SpCas9-NG achieved a high editing efficiency for 10 out of 12 TAG stop codons when swapping to TAA (**Figure S8B**). We then swapped the two remaining TAGs by a *URA3* in-and-out strategy, similar to the one described for the *ENA* genes. A detailed table summarizes the correction steps made to each version of *synIV* (**Table S4**). Whole genome sequencing revealed that the final *synIV* strain (yeast_chr04_9_07) has several variants from the original design, all of which are listed in **Table S5**.

The fitness of yeast_chr04_9_07 is near wild type under various conditions (**Figure 3C**, **Figure S8C**), indicating that the *in silico* designed *synIV* is capable of powering a yeast cell to near normal growth rates. It is well documented that the yeast genome is well-organized within the three-dimensional nuclear space^29^. Consequently, we wondered whether the sequence changes made to chromosome IV could affect the 3D structure of *synIV*. We performed Hi-C on both wild-type and *synIV* strains, and as expected due to the lack of Ty element repeats, we found the intra-chromosomal contact map of *synIV* was almost completely free of unmappable regions, which appear as white stripes on the wild-type *chrIV* map (**Figure 3D**). Notably, in *synIV* we found increased interactions between *Ty*-flanking sequences reflecting the fact that the intervening *Ty* elements were removed during synthesis. Furthermore, the 3D projections of the Hi-C maps showed both *synIV* and wild-type IV chromosomes are organized similarly within the intranuclear space (**Figure 3E**, supplementary movies), suggesting that Sc2.0 modifications have only minor effects on 3D structure of synthetic DNA molecules, but increase the contact map mappability of Hi-C.

Transcript profiling revealed a small number of genes that were differentially expressed in the *synIV* strain (**Figure 3F**), which encode uncharacterized open reading frames (ORFs) according to the *Saccharomyces* gene database (SGD). Briefly, for upregulated genes in *synIV*, *YDL007C-A*’s promoter is adjacent to a serine tRNA gene, and *YDR278C* overlaps with a glutamate tRNA gene in wild-type IV. It is well known that tRNAs silence nearby polymerase II transcription^30,31^, so we speculate the upregulation of *YDL007C-A* and *YDR278C* are caused by the removal of the flanking tRNA genes. For down-regulated genes in *synIV*, *YDR541C* is now adjacent to the new universal telomere cap (UTC) after deleting the subtelomeric region in *synIV*, so the down-regulation of *YDR541C* is likely due to telomere silencing.

Given the 479 identical loxPsym sites incorporated into *synIV*, we wondered whether *synIV* was prone to chromosome rearrangement or loss during prolonged culturing. We generated a heterozygous diploid *synIV* strain, with a *URA3* gene inserted on the left arm of either *synIV* or *wtIV*, and then inserted a *KanMX4* marker on the same allele as *URA3* (**Figure S9A**). We passaged both diploid strains for approximately of 100 generations, and plated the cells on YPD and 5-FOA medium for colony counting. We noticed a higher fraction of colonies were 5-FOA resistant when the *URA3* gene was on *synIV* (**Figure S9B**). PCR examination showed all the 5-FOA resistant colonies had completely lost the *URA3* gene (**Figure S9C**). Also, 100% of the 5-FOA resistant colonies were G418 resistant (**Figure S9D**), indicating that the loss of *URA3* was due to mitotic recombination and not chromosome loss.

### “Inside out” *synIV* reveals limited gene expression changes

Yeast chromosomes are organized in a dynamic nonrandom architecture dominated by a set of contact points with the spindle pole body and the inner nuclear membrane in the nucleus^29^, and such organization influences gene regulation. Here, we wondered whether the 3D architecture of *synIV* is sensitive to manipulation or whether it can be synthetically redefined in a living yeast cell. As a proof of concept, we sought to flip the linear orientation of chromosome arms. To accomplish this, we relocated the most structurally impactful sequence, *CEN4*, to a context as far-removed from its native location as possible, namely the telomere-adjacent sequence, thus moving the telomere-adjacent sequence towards the centromere. The goal was to create an “inside out” *synIV* chromosome, assuming that the newly relocated *CEN4* remains connected to the spindle pole body (**Figure 4A**). This move places the former centromere proximal sequences adjacent to the (former) subtelomeres, and the former telomere-proximal sequences adjacent to the centromere. We refer to this as an “inside out” (IO) chromosome.

**Fig. 4.**
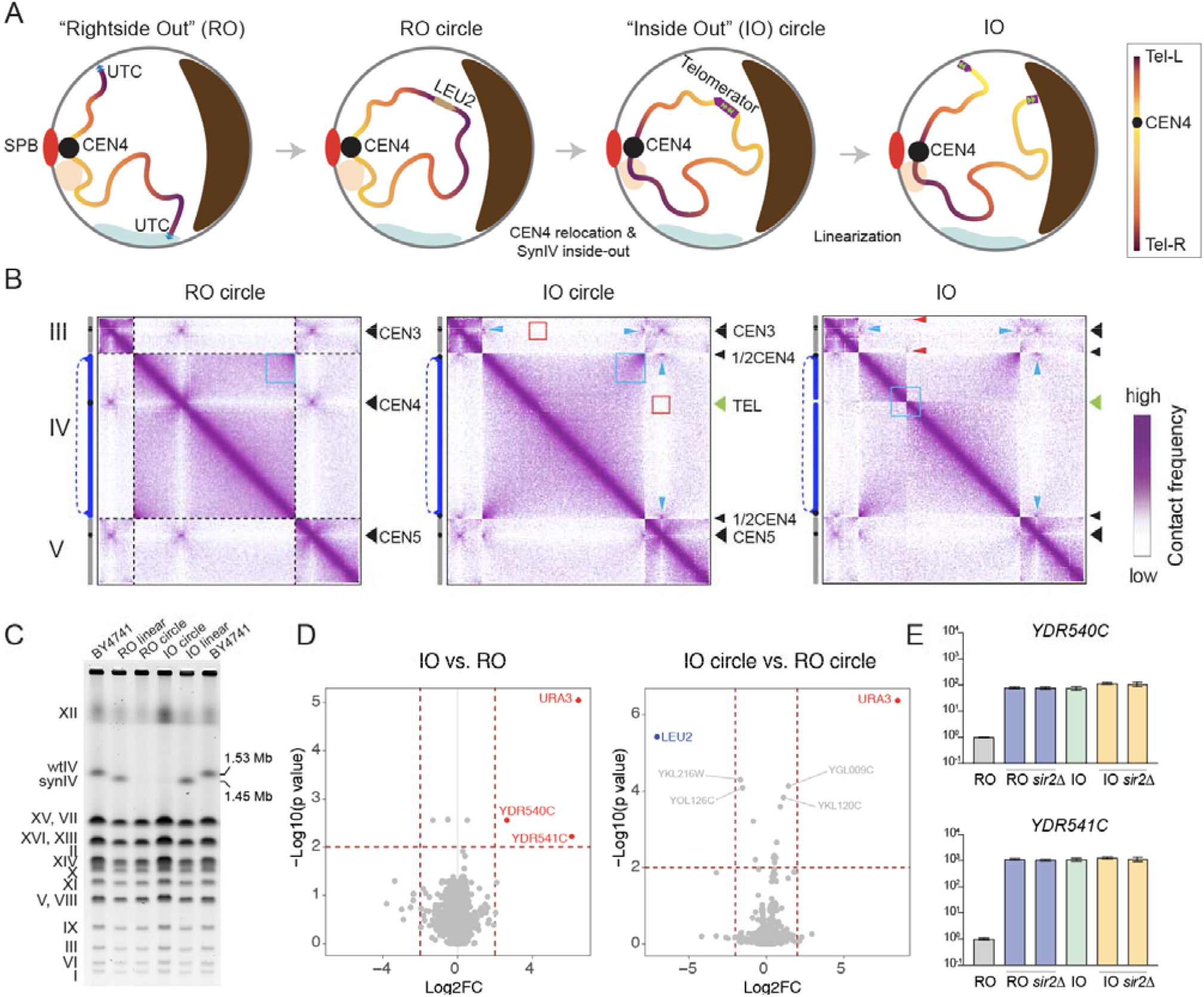
“Inside out” *synIV* engineering and characterization. (**A**) Schematic illustration of the “inside out” *synIV* construction. *SynIV* chromosome is shown as a gradient color line, with chromosome regions near the native centromere shown by light yellow and chromosome regions near the native telomeres shown by dark purple. Two hypothetical sub nuclear compartments are shown as orange and light green regions near the nuclear envelope. The nucleolus is depicted in brown. SPB, spindle pole body. (**B**) Hi-C contact maps of yeast with *synIV* in different configurations annotated in blue on the left side of each map (5 kb-binned). Black and green arrowheads indicate centromere and telomere positions, respectively. Violet to white color scale reflects high to low contact frequency (log10). Blue boxes, red boxes, blue arrows, red arrows indicate the contact map changes upon structural manipulation (see main text). (**C**) Linear *synIV*, circular *synIV* and linear wild-type IV were separated by pulsed field gel electrophoresis. CHEF mapper auto algorithm was used for separating the chromosomes, low molecular weight, 200 kb; high molecular weight, 2500 kb; voltage, 6 V/cm; total electrophoresis time was 29 hours. (**D**) Transcript profiling of both linear and circular *synIV*s. *URA3* marker was present in IO circle *synIV*, *LEU2* marker was present in RO circle *synIV*. Fold change cut off is 4, p-value cutoff is 0.01. (**E**) *YDR540C* and *YDR541C* mRNA levels were determined by RT-qPCR in the indicated *synIV* strains with or without *sir2* deletion. Bars represent mean ± SD of three technical replicates.

To construct the inside out *synIV* strain, we first circularized the original “rightside out” (RO) *synIV* by providing a *LEU2* marker with flanking homology arms. The left homology arm maps to the last chunk of *synIV* (chunk JJ5) whereas the right homology arm maps to the first chunk of *synIV* (chunk A1). DNA double-strand breaks were introduced by CRISPR-Cas9 near both UTCs, resulting in their excision and the simultaneous bridging of the sequence by the *LEU2* donor DNA (**Figure 4A, Figure S10A**). Next, we performed intramolecular centromere relocation by simultaneously replacing *LEU2* with *CEN4* and replacing *CEN4* with the telomerator, a synthetic cassette designed to convert circular DNA to linear DNA *in vivo*^32^ (**Figures S10B-C**). We expected that the relocated *CEN4* sequence would retain contact with the SPB, forcing the original telomere-proximal sequences close to the SPB and flipping the orientation of both *synIV* chromosome arms. Lastly, we transiently expressed I-SceI endonuclease in the IO circle *synIV* strain to linearize the IO circle at the telomerator (**Figure 4A, Figure S10D**). Hi-C maps were generated to validate these extraordinary structural reorganizations of *synIV*. Precisely as we predicted, 1) telomere fusion resulted in the formation of a circular RO chromosome, whose formerly peri-telomeric ends gave rise to strong intra-chromosomal contacts (**Figure 4B**, first panel blue box), while their interactions with the remaining active telomeres were lost. 2) After *CEN4* relocation, the interaction between former *CEN4* with other CENs was now absent (**Figure 4B**, second panel red boxes), instead the ectopic relocation of *CEN4* within the former peri-telomeric region contact strongly with the other two centromere (**Figure 4B**, second panel blue arrows), indicating the relocation of CEN4 is sufficient to flip the entire circular *synIV* inside out. 3) Lastly, the re-linearization of the inside out circle *synIV* at the former peri-centromere gave rise to an inside out linear *synIV*, and the new *TEL* sequences appeared interacting with other telomeres (**Figure 4B**, third panel red arrows). Pulsed field gel electrophoresis was performed for the two linear *synIV* strains (RO and IO) and two circular *synIV* strains (RO circle and IO circle). Both RO and IO *synIV*s are slightly smaller than wild-type IV in size, as we deleted 77.3 kb in *synIV* compared to the wild type IV. Both RO circle and IO circle *synIV*s are trapped in the agarose plug, as circular DNAs are unable to migrate out of the plugs through PFGE^33^ (**Figure 4C**).

With these structurally engineered *synIV* strains, we asked whether their forced chromosome-wide gene repositioning in the IO *synIV* led to specific gene expression changes. Surprisingly, only two telomeric genes, *YDR540C* and *YDR541C,* were upregulated in the IO linear vs. RO linear *synIV* comparison (**Figure 4D**, left). The IO circle vs RO circle did not show any differentially expressed genes on entire chromosome IV (**Figure 4D**, right). The two telomeric genes were immediately adjacent to the RO *synIV* right telomere, and were relocated to the centromere cluster near the SPB in the IO *synIV*, far away from the telomere. This indicates that relocation of *CEN4* brought these two genes from a telomeric repression domain to a more active domain. Further, the silencing of *YDR540C* and *YDR541C* in the RO *synIV* was SIR protein dependent, likely through a histone deacetylation mechanism as previously reported for other telomeric genes^34,35^. Knocking out *sir2* in RO *synIV* completely abolished the silencing effect (**Figure 4E**). To test whether genes on other chromosomes have similar minor expression changes upon repositioning, we compared the transcriptome profiles of two *CEN8* relocation strains (one with *CEN8* relocated to the extreme left end of the chromosome, the other was an isochromosome *VIII* with *CEN8* relocated at the extreme right end of the chromosome, **Figure S11A**, described in detail in Lauer et al.^36^) with the wild-type strain. Besides effects on a few dubious ORFs on other chromosomes, we found that gene expression was largely unchanged in both strains when compared to native *VIII* (**Figures S11B-D**). Together, our results suggest that relative gene positions in the interior space of the nucleus have only minor effects on gene expression, in agreement with the hypothesis that the major nuclear compartments regulating gene expression in yeast are 1) the compartment near nuclear envelope^19^ and 2) the compartment near nuclear pore complexes^37^.

### Transcriptional repression of *synIV* by multipoint tethering to inner nuclear membrane

The ability to control genome organization allows us to interrogate the relationship between gene position and gene regulation, and ultimately gain new insights on cellular functions. In mammalian cells, recruiting a single specific genomic locus to the nuclear periphery led to modest genome repression by hijacking interactions between LacI and LacO^38^, or dCas9/gRNA and targeted genomic loci^39^. However, whole chromosome repression is difficult to obtain, due to the lack of a pervasive distribution of targetable sequences in a chromosome of interest. The Sc2.0 genome incorporates near 4000 identical loxPsym sites to the downstream of nonessential genes^5^, serving as the ideal genome for testing the possibily of whole yeast chromosome/genome repression. Thus, we sought to tether *synIV*, the largest synthetic yeast chromosome, to the inner nuclear membrane (INM) at up to 479 positions, and asked whether whole chromosome repression could be achieved. We first employed ZFDdesign^40^, a machine learning platform for zinc finger protein design, and generated two loxPsym site binding zinc finger proteins, one (ZF-SAB) skips one nucleotide in every six nucleotides, the other one (ZF-Ext) does not incorporate nucleotide skipping (**Figure S12A**). We tested the binding activity of both zinc finger proteins to loxPsym using a bacterial one-hybrid assay^41^, and we found that ZF-Ext bound 18 bp of loxPsym strongly (**Figure S12B**). We then anchored ZF-Ext (hereafter referred to as “ZF”) to the INM by fusing it to the C-termini of two INM-localized proteins, HEH1 and HEH2^42^. To achieve dynamic control of *synIV* tethering, we expressed ZF fusions from an inducible promoter (ZEVpro) containing 6 tandem binding sites for mouse transcriptional factor Zif268/Erg1, fused to the estrogen binding domain and the VP16 activation domain^43^ (**Figure 5A**). Due to potential overexpression effects of HEH1 and HEH2 themselves, we designed inducible HEH1/ HEH2-only constructs as controls. BY4741 and *synIV* transformed with HEH1/HEH2-only, or HEH1-ZF/HEH2-ZF were grown in medium with estradiol for growth curve measurements. *synIV* had a much stronger growth defect than BY4741 when HEH1 and HEH2 were fused with the ZF (**Figure 5B**). The slight growth defect of BY4741 when ZF is present is attributed to weak non-specific binding of ZF to the wild-type genome. Singly expressed HEH1-ZF or HEH2-ZF fusions were insufficient to inhibit growth of *synIV* (**Figure S12C**). To determine whether the growth defect was a consequence of *synIV* repression, we performed RNA-seq for BY4741 and *synIV* strains with ZF fused to HEH1/HEH2 (or not), and counted differentially expressed genes on chromosome IV. Unlike BY4741, which had similar numbers of downregulated and upregulated genes (81 vs 85), we found many more downregulated genes (302) in *synIV* vs. 40 downregulated (**Figure 5C**). Further, the downregulated genes from BY4741 mostly overlapped the set of genes downregulated in *synIV* (**Figure 5D**). The multipoint tethering mediated repression appeared to have no bias toward gene locations on the chromosome (**Figure 5E**), nor gene essentiality (**Figure S12D**), however, the degree of repression positively correlated with the distance between the start codon and the nearest loxPsym site (**Figure 5F**). We also measured the distances between the telomere and the first repressed gene’s loxPsym on each chromosome arm (**Figure S12E**), depicting the size of the telomeric repression domain. Lastly, to rule out the possibility that whole chromosome repression was caused by excessive ZF binding on *synIV*, we constructed ZF only controls and did not observe growth defects (**Figure S12F**).

**Fig. 5.**
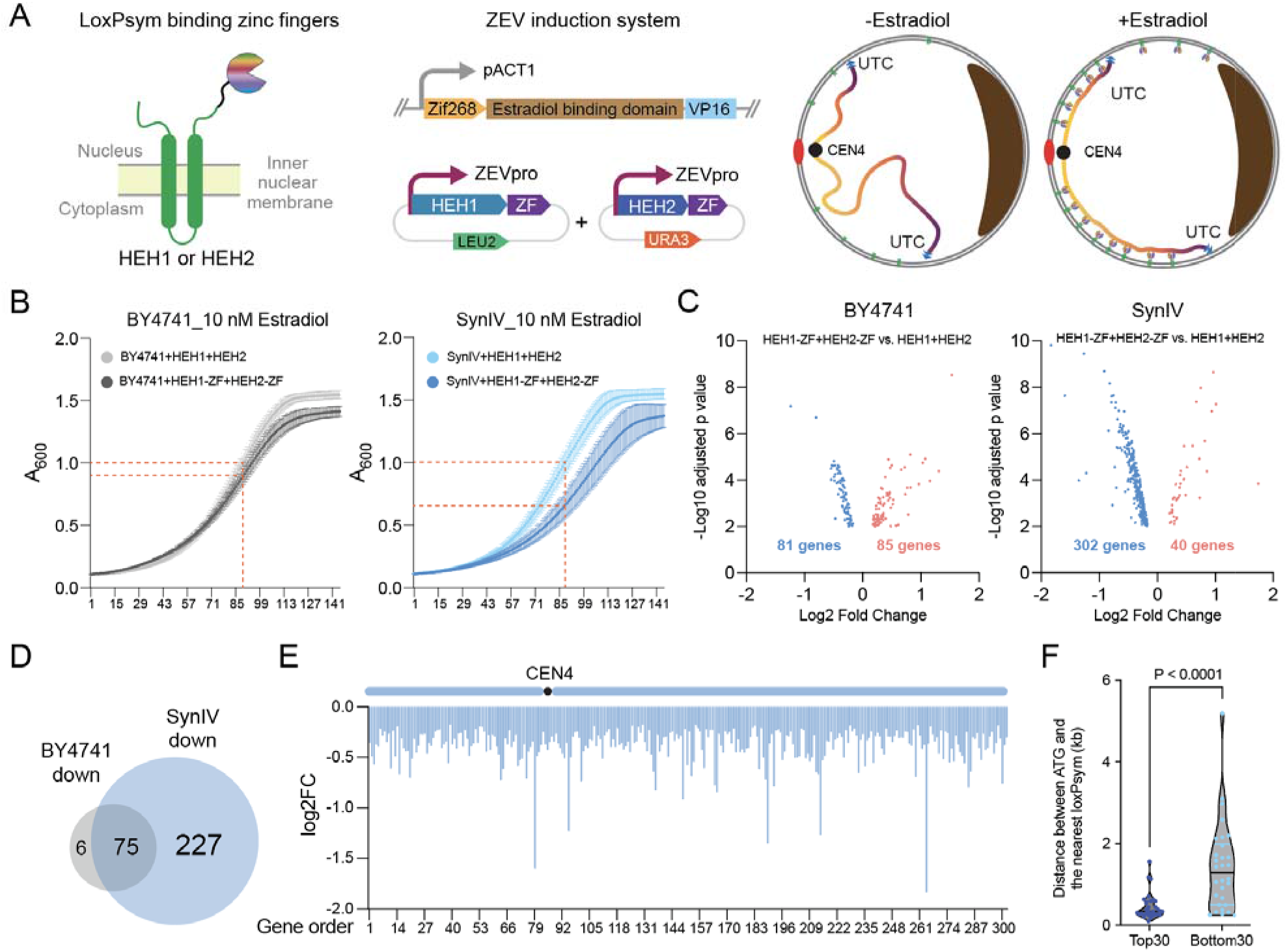
*SynIV* tethering to the inner nuclear membrane. (**A**) Schematic illustration of *synIV* multipoint tethering assay. Zinc finger protein was fused to both *HEH1* and *HEH2* under an estradiol-inducible promoter (ZEVpro). Mouse *Zif268* gene fused with estrogen binding domain and VP16 activation domain^43^ was expressed under the *ACT1* promoter and the whole cassette was integrated in the *YKL162C-A* locus on chromosome *XI*. In the presence of estradiol, HEH1-ZF and HEH2-ZF will be anchored to the INM allowing multipoint (up to 479 positions) tethering the loxPsym containing yeast chromosome to the nuclear envelope. (**B**) Growth curve of BY4741 and *synIV* strains contain either HEH1/HEH2 only, or HEH1-ZF/HEH2-ZF constructs. A_600_ was measured in every 10 min, X axis is the A600 reading points, Y axis is A_600_. Yeast cells were harvested at the timepoint at which HEH1/HEH2 containing strains reached to A_600_=1. Bars represent mean ± SD of six technical replicates. (**C**) Volcano plots showing differentially expressed chromosome IV genes in BY4741 (left) and *synIV* (right). Adjusted p value cutoff was 0.01. (**D**) Venn diagram showing that downregulated genes in BY4741 and *synIV* largely overlapped. (**E**) Bar plot showing *synIV* downregulated gene distribution, X axis is the order of downregulated gene from left arm to right arm, Y axis is fold change. (**F**) Distances between ATG and the nearest loxPsym site for the top 30 and bottom 30 genes on the *synIV* downregulated gene list. P value was analyzed using an unpaired two-tailed t test.

### Selectively repressing loxPsym containing chromosome or chromosome segment

To demonstrate the generalizability of synthetic chromosome multipoint tethering mediated repression, we introduced the tethering system into *synIII* strain, as well as a semi-*synIV* strain that has only a ∼500 kb synthetic region (megachunk X-JJ). Evident growth defects were observed in the two synthetic strains but not in BY4741 when ZF was fused to HEH1 and HEH2 (**Figure 6A**). Transcriptomic analysis showed that *synIII*, and *synIV X-JJ* (but not the megachunk A-W region on the same chromosome) were specifically repressed after tethering them to the INM (**Figure 6B, Figure S13**). Since multipoint tethering mediated chromosome repression is induced by estradiol, a chemical that can be easily removed from the culture medium, tethering-mediated chromosome repression should be reversible. Native, tethered and untethered *s* and *synIV X-JJ* were harvested for both growth assay and RT-qPCR analysis. Tethering-*nIII* associated growth defects were indeed reversed in the untethered condition (**Figure 6C**). Also, downregulation of a representative gene on *synIII* or *synIV X-JJ* were restored upon untethering (**Figure 6D**). Together, synthetic chromosome tethering is generalizable for repression of whole syn-chromosomes or specific syn-chromosomal regions, and such repression can be easily lifted.

**Fig. 6.**
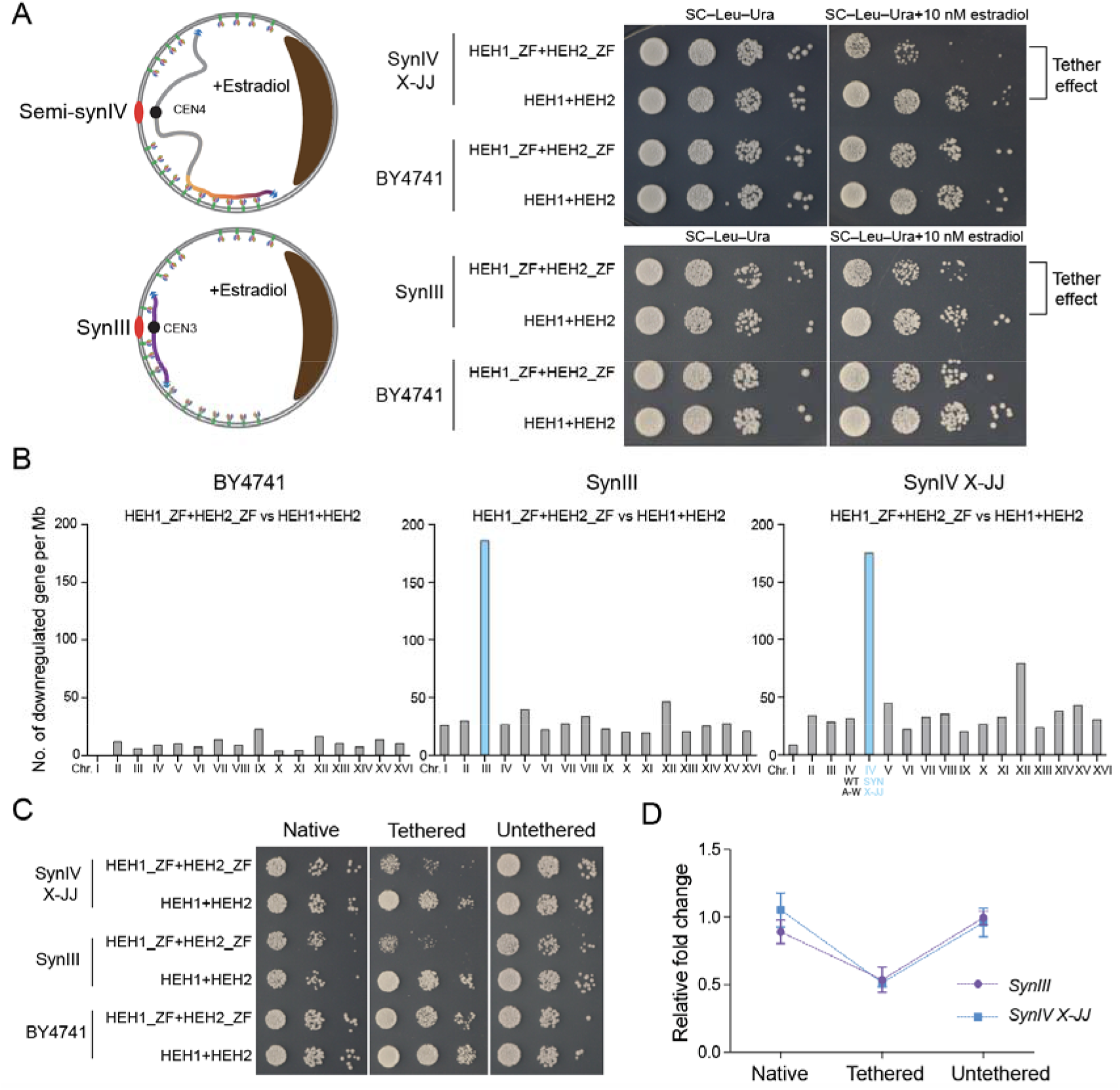
*SynIII*, semi-*synIV* multipoint tethering. (**A**) Semi-*synIV* and *synIII* tethering schematic and 10-fold serial dilution spot assay on SC–Leu–Ura medium or SC–Leu–Ura medium supplemented with 10 nM estradiol. (**B**) Downregulated gene counts in BY4741, *synIII* and *synIV X-JJ* strains in the context of chromosome tethering. X axis is chromosome or chromosome segments, Y axis is number of downregulated gene per megabase. (**C**) Semi-*synIV* and *synIII* reversible tethering 10-fold serial dilution spot assay on SC–Leu–Ura medium (left and right panels) or SC–Leu–Ura medium supplemented with 10 nM estradiol (middle panel). (**D**) RT-qPCR analysis of *YCR020W-B* on *synIII*, and *YDR321W* on *synIV* in native, tethered and untethered conditions. Fold changes were normalized to the HEH1+HEH2 only controls in each strain. Bars represent mean ± SD of three technical qPCR replicates.

## Discussion

The assembly of *synIV*, the largest synthetic yeast chromosome, is a key step towards the completion of the first synthetic eukaryotic genome. In this study, we deployed the “design-build-test-learn” cycle and acquired new insights on eukaryotic genome design, assembly methodology, debugging strategy and importantly, 3D genome organization-transcriptional regulation relationships. *SynIV* by itself is a valuable resource for studying fundamental questions regarding genome evolution, chromosome stability. Also, the fairly evenly distributed 479 loxPsym sites on *synIV* is a unique and novel resource to study 3D organization of synthetic chromosomes. The nuclear envelope tethering assay demonstrated here exemplifies a new concept: programmable whole chromosome silencing. During the transition from steady state to the silenced state, nucleosome occupancy changes, histone modifications such as H4K16 acetylation changes might be worth examining. Given that the tethering is specific to the synthetic portion of a semisynthetic chromosome (**Figure 6**), a library of semi-synthetic strains can be easily generated via meiotic recombination from heterozygous diploid synthetic strains, so the loxPsym site containing “patches” can be selected based on the most interesting distributions. Furthermore, these loxPsym sites enable Cre-mediated synthetic chromosome shuffling (SCRaMbLE^44–54^), a powerful application of synthetic chromosomes for novel genomic function discovery.

Proper positioning of synthetic chromosomes in the yeast nucleus ensures the physiological expression of genes on the chromosome. How to define the organization of a synthetic chromosome is a worthwhile question. *SynIV* carries the most genes in *S. cerevisiae*, changing *synIV*’s organization should be most impactful in terms of changing gene-gene interaction, and relative 3D positions. Here, we manipulated *synIV*’s 3D organization by design, effectively turning it inside out, representing a major shift from sequence rewriting to structural redefinition of a synthetic yeast chromosome. Mammalian chromatin is packaged into topological association domains (TADs) in mammalian nucleus and plays important roles in regulating gene spatiotemporal expression^55^. In contrast, yeast sub-nuclear compartments are seemingly restricted to the part adjacent to the nuclear envelope. Whether yeast has mammalian TAD-like territories that reside in the interior space of the nucleus is unclear^56–58^. Through the construction and characterization of the “inside out” *synIV* strain, we made drastic spatial changes to all the genes on *synIV*, and we found only minor gene expression changes, leading to the conclusion that specific positioning in the *interior* space of yeast nucleus plays a negligible role in gene expression regulation. The inside out *synIV* also changes the relative gene positions between *synIV* genes and other wild-type genes. However, these changes overall did not have substantial effects on gene expression, suggesting that gene-gene interaction plays a negligible regulatory role in yeast. Our results support the hypothesis that local self-associating domains are the major chromosomal interaction forms that affect transcriptional regulation^56^, most regulation is local rather than regulated by global intranuclear position in the nucleoplasm.

So far, we performed multipoint synthetic chromosome tethering assays in three different strains, and the numbers of downregulated (at an FDR<0.01) genes were consistently around 30% of the total gene numbers in those synthetic regions (31.1% for *synIII*, 27.8% for *synIV* full length and 30.5% for *synIV X-JJ*). This may be attributable to the somewhat weak binding activity of the zinc finger protein to the loxPsym sequence (**Figure S12A-B**). Designing more zinc finger proteins with different binding affinities may help establish stronger or milder silencing of synthetic chromosomes in the future.

Live cell imaging is a powerful tool to study dynamics of chromosome conformational changes. Previous efforts focused on a few genomic loci due to targeting limitations^59,60^. Taking advantage of the near evenly spaced loxPsym sites on *synIV*, one can fuse a fluorescence protein or split forms thereof to the loxPsym binding zinc finger protein, and monitor architectural changes during cell divisions in real time.

In summary, we showed that *synIV*, a megabase-sized eukaryotic chromosome can be *in silico* designed and *de novo* synthesized, and its 3D organization can be synthetically reprogrammed for probing genome organization-transcriptional regulation relationships. Thanks to the hundreds of identical loxPsym sites incorporated, this method opens up many chromosome architectural manipulation possibilities for in-depth understanding of the genome dynamics using unique synthetic chromosomes like *synIV*.

### Limitations of the Study

In our study, we reached the conclusion that *synIV* has a higher mitotic recombination frequency than the wild-type *chrIV*. However, we have not investigated whether this increase is directly linked to any specific feature(s) of *synIV*. Considering that *synIV* represents an extreme case as being the largest synthetic yeast chromosome, further tests on the mitotic recombination frequencies of other Sc2.0 chromosomes with smaller and intermediate sizes could be informative. Also, understanding whether the location of crossovers among different recombinants is consistent or random could provide insights for the elevated mitotic recombination frequency of *synIV*. Lastly, for the multi-point synthetic chromosome tethering assay, direct visualization approaches such as live cell imaging, or fluorescence *in situ* hybridization (FISH) could be implemented to further tie the silencing effects to the 3D genome organization changes and provide direct visualization of the relocalized chromosome, potentially shedding light on the actual number of tethering points in a given cell.

## Supporting information

3D projection

## Acknowledgements and competing interests

We thank the NSF for grants MCB-1026068, MCB-1443299, MCB-1616111 and MCB-1921641 to JDB for supporting this work. This work is also supported by National Natural Science Foundation of China (32150025, 32030004, 31725002), Shenzhen Science and Technology Program (KQTD20180413181837372), Shenzhen Outstanding Talents Training Fund and Bureau of International Cooperation, Chinese Academy of Sciences (172644KYSB20180022) to JD. This work was supported by the UKRI EPSRC Fellowship (EP/V033794/1) to G.S. The synthesis of the left arm starting materials [proposal award DOI “Community Science Program 1110” (CSP 1110) titled “Synthesis of the largest yeast chromosome, chromosome IV, and the synthetic yeast genome Sc2.0” was conducted by the U.S. Department of Energy Joint Genome Institute (https://ror.org/04xm1d337), a DOE Office of Science User Facility, was supported by the Office of Science of the U.S. Department of Energy operated under Contract No. DE-AC02-05CH11231. We thank Meghan O’Keefe for collecting Build-A-Genome authors’ information. We thank André M. Ribeiro-dos-Santos’s help with depositing the data to the Gene Expression Omnibus.

This work is part of the international Synthetic Yeast Genome (Sc2.0) consortium. The chromosome design and building consortium includes research groups worldwide: Boeke Lab at Johns Hopkins University and New York University (led chromosomes I, III, IV, VI, VIII, IX), Chandrasegaran lab at Johns Hopkins (led chromosomes III and IX), Cai Lab at University of Edinburgh and University of Manchester (led chromosomes II, VII and tRNA neochromosome), Yue Shen’s team at BGI-Research Shenzhen (led chromosomes II, VII, XIII), Y.J. Yuan’s team at Tianjin University (led chromosome V, X), Dai Lab at Tsinghua University and Shenzhen Institute of Advanced Technology, CAS (led chromosome XII), Ellis Lab at Imperial College London (led chromosome XI), Sakkie Pretorius’s team at Macquarie University (led chromosomes XIV, XVI), Matthew Wook Chang’s team at National University of Singapore (led chromosome XV), Bader and Boeke Labs at Johns Hopkins University (led design and workflow), and Build-A-Genome undergraduate teams at Johns Hopkins University and Loyola University Maryland (contributed to chromosomes I, III, IV, VIII, IX). The Sc2.0 consortium includes numerous other participants and are acknowledged on the project web site www.syntheticyeast.org.

## Author Contributions

J.D.B., W.Z., J.S.B. L.A.M. conceptualized the study. J.D.B., J.D., Y.A., S.D., L.S., K.Z. supervised the study. W.Z., L.L.S, H.Y., M.J.S., H.K., E.V., M.A.B.H., X.S., Q.J., S.L.L., L.H.M, Y.Z., N.E., S.J.L., B.R.C., Y.Z., Z.X., M.S. performed the experiments, including chunk DNA assembly, megachunk DNA assembly, *synIV* intermediate strains integrations, Hi-C library construction and data processing, RNA sequencing analysis. J.D.B., L.A.M., G.S., J.S.B. designed *synIV*. V.F., G.S. performed the whole genome analyses. M.B.N., G.W.G., D.M.I. designed and tested the loxPsym binding zinc finger proteins. Build-A-Genome class students contributed the chunk DNA assembly. J.C. and J.S.B. uploaded the *synIV* sequencing datasets to NCBI. Manuscript was drafted by W.Z., reviewed and edited by L.L.S., M.J.S., M.A.B.H., Y.A., J.D. and J.D.B.

## STAR Methods

### Key resources table

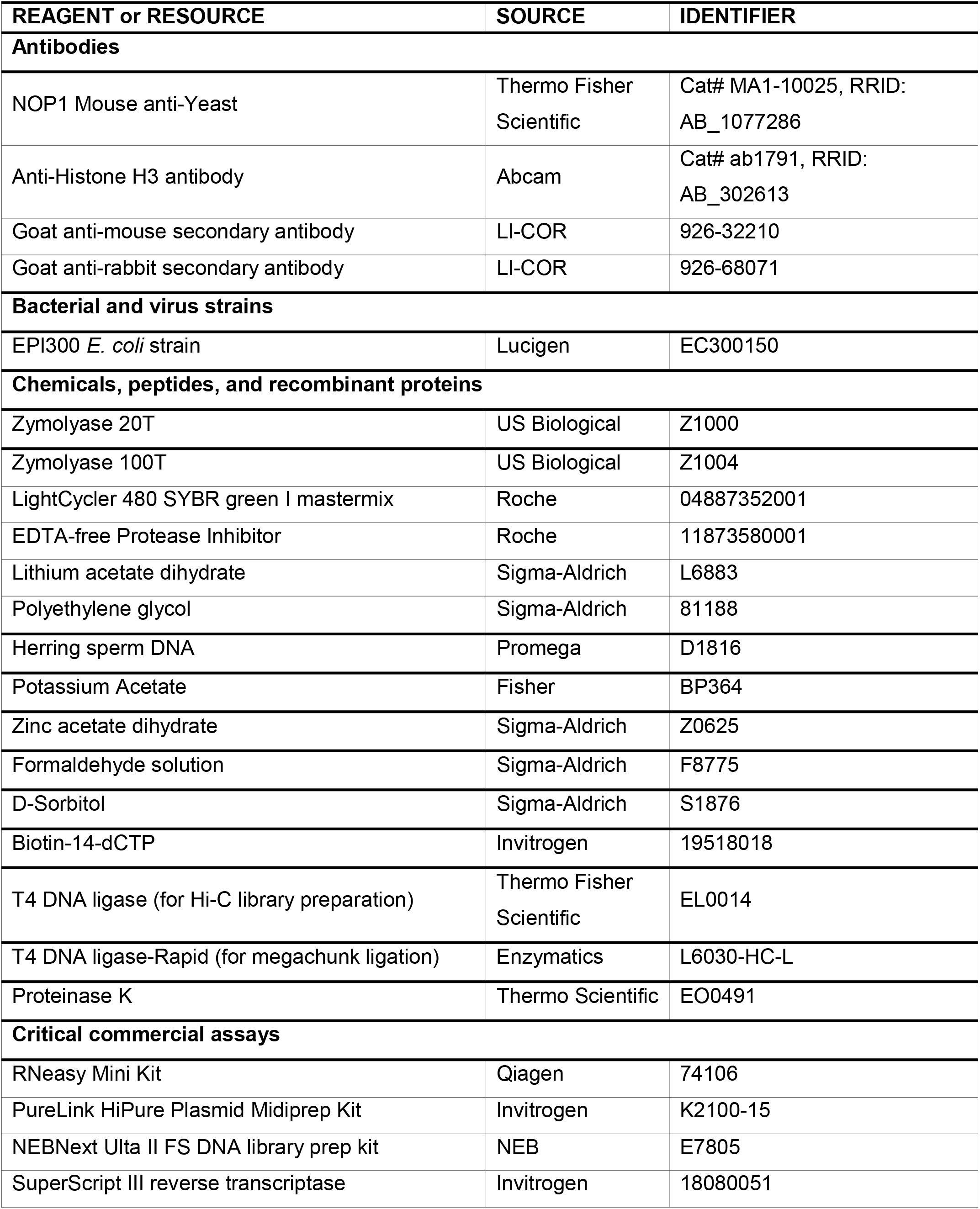
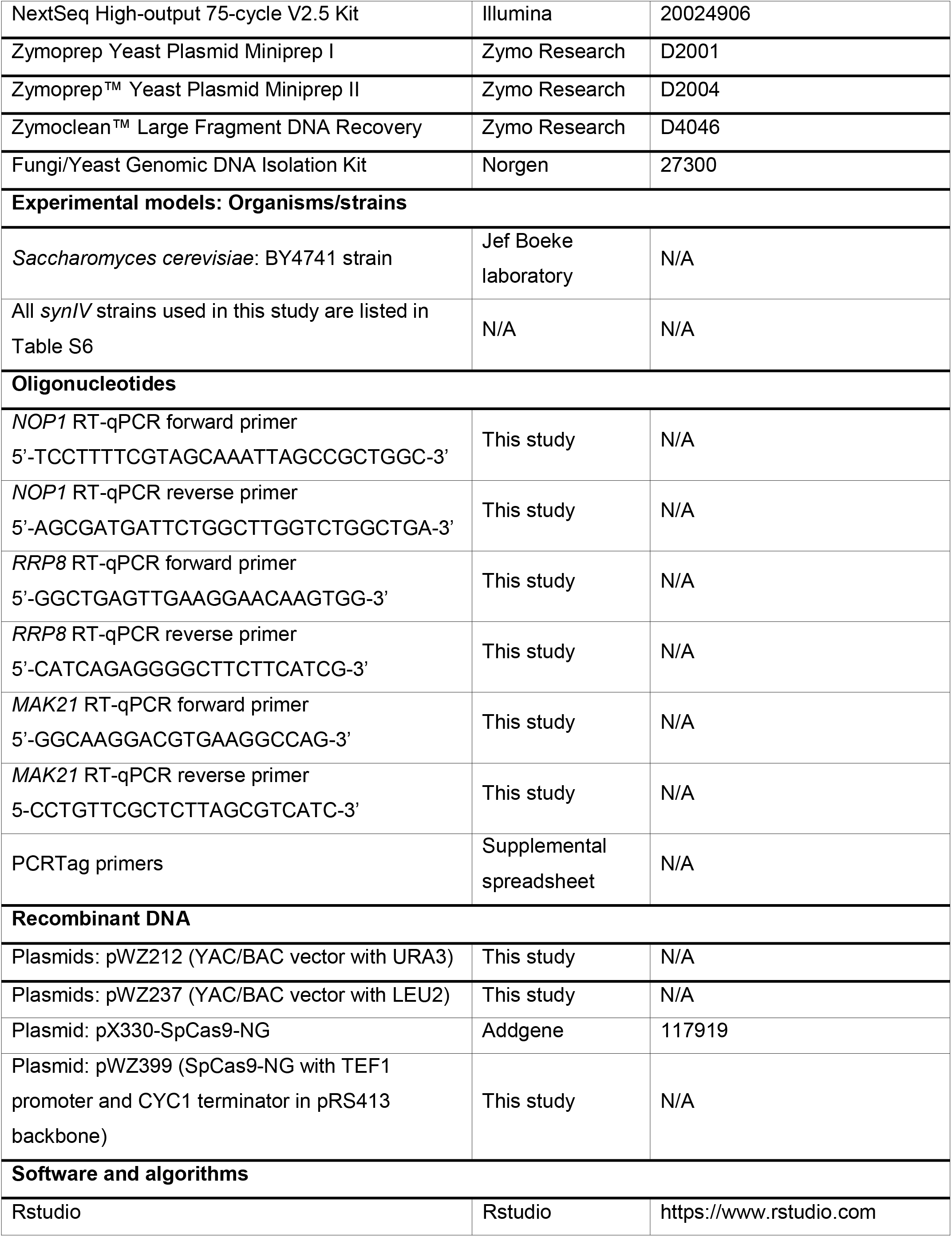
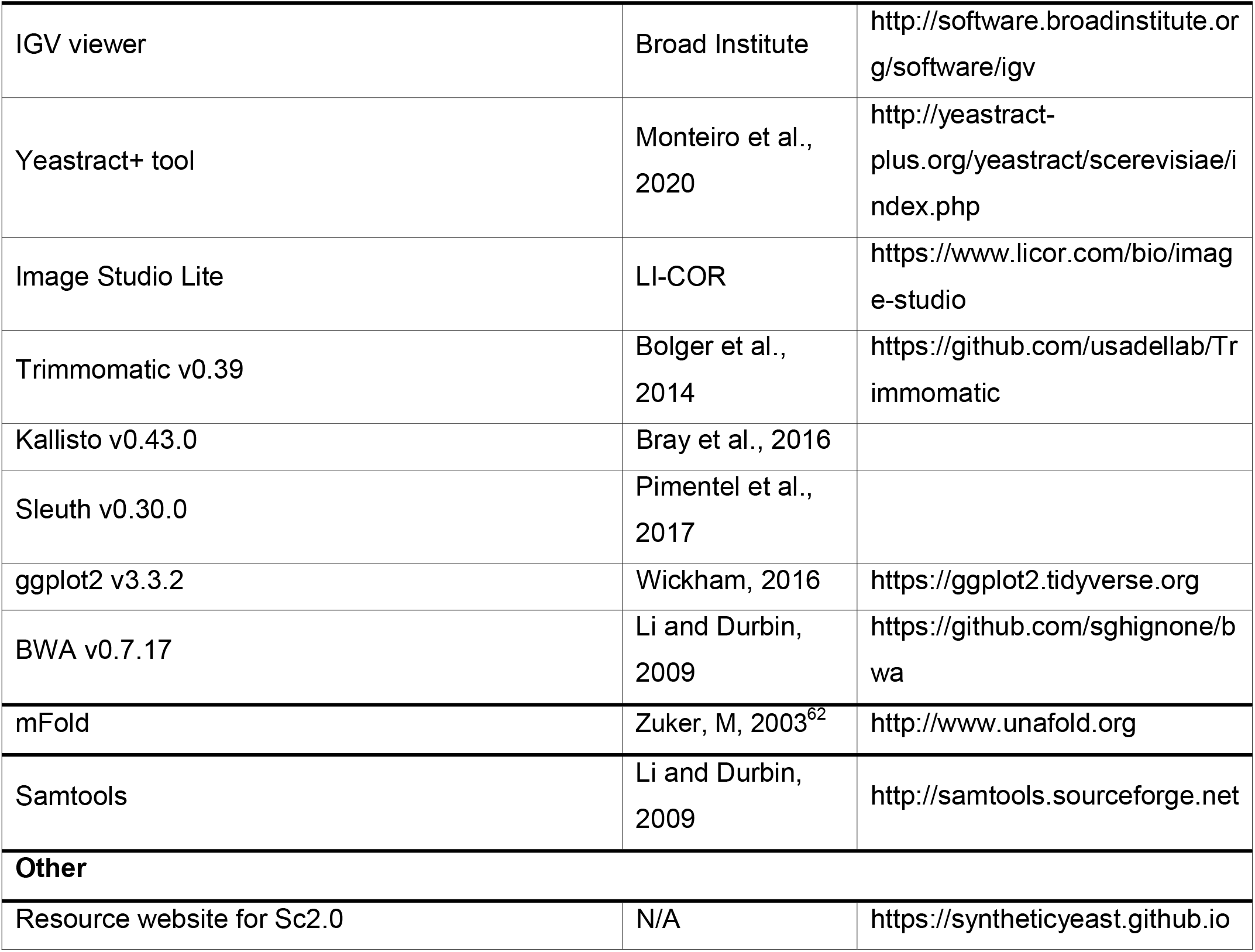

#### RESOURCES AVAILABILITY

##### Lead contact

Further information should be directed to and will be fulfilled by the lead contact Jef D. Boeke

##### Materials availability

All unique/stable reagents generated in this study are available from the Lead Contact with a completed Materials Transfer Agreement.

##### Data and code availability

Data: All data reported in this paper have been deposited to Gene Expression Omnibus accession number is GSE239536 (subseries GSE239532 for Hi-C, subseries GSE239534 for RNA-seq, subseries GSE239535 for WGS).

Code: This work did not generate any new code.

Any additional information required to reanalyze the data reported in this paper is available from the lead contact upon request.

### Experimental Methods

#### Vectors, strains, and media

YAC-BAC shuttle vectors pWZ212 and pWZ237 were constructed by former Boeke lab member, Leslie Mitchell. Parental strains for intermediate *synIV* entry were from the yeast knock-out collection. All *synIV* strains used in this study are listed in Table S6. YPD, YPG and synthetic complete (SC) media were prepared following standard recipes. Media with DNA damage reagents, DNA replication inhibitors, high pH, oxidative stress, high osmotic pressure etc. were prepped freshly by following previous publications^7,9,13^.

#### Chunk assembly

Three or four “minichunks” were either released from their vector backbones by restriction enzyme digestion, or amplified by PCR. Approximately 200 ng of each minichunk fragment, ∼10 ng linearized pRS413 acceptor vector and ∼50 ng of each linker fragment were co-transformed into yeast. Yeast cells were plated on SC–His medium for 3 days of growth at 30°C. Junction PCR was performed to identify yeast clones that contain correctly assembled chunks. Chunk DNA was isolated from yeast using a yeast plasmid miniprep kit (Zymo research, D2004), and further transformed into *E. coli* for amplification. Correctly assembled chunks were verified by PacBio sequencing.

#### Megachunk assembly, cloning and integration

Megachunks were assembled from three to six “chunks”, each previously released from its vector backbone via rare cutting restriction enzyme digestion (Table S2). Following gel extraction of the chunk fragments, chunk fragments were pooled and ligated at 16°C with the following components: ∼200 ng of each gel-purified chunk DNA fragment, 1 μL of T4 DNA ligase buffer, 0.5 μL of T4 DNA ligase (Enzymatics, L6030-HC-L) and H_2_O to a final volume of 10 μL. The ligation product was transformed into yeast with a YAC (yeast artificial chromosome)-BAC (bacterial artificial chromosome) shuttle vector and two linker DNAs. The YAC-BAC vector was either pWZ212 (with *URA3* marker for megachunks that contain *LEU2* marker) and pWZ237 (with *LEU2* marker for megachunks that contain *URA3* marker) (Fig. S1). Yeast cells were plated on SC–Leu–Ura dropout agar medium for three days of growth at 30°C. Yeast colonies were verified by junction PCR for all chunk fragments. Either forward or reverse of a primer pair must be chosen from synthetic PCRTag to distinguish megachunk junctions from native yeast genome.

Megachunk plasmids (BACs) were recovered from yeast by using a yeast plasmid miniprep kit (Zymo Research, D2001), and subsequently electroporated into the EPI300 *E. coli* strain (Lucigen, EC300150). Megachunk plasmids were extracted from EPI300 *E. coli* strain by using PureLink™ HiPure Plasmid Midiprep Kit (Invitrogen, K2100-15). Megachunks were excised using restriction sites designed at left and right ends (**Table S3**) and transformed into BY4741 or a derivative, with selection for Leu or Ura markers as appropriate, and replica plated to verify loss of the resident marker to perform “SwAP-In” (Dymond et al., 2011, Richardson et al. 2017). The intermediate strains, as well as the versions of *synIV* and the strains containing them are described in Table S6.

#### PCRTagging

Genomic DNA was prepared as follows. Yeast colonies were scraped from agar medium and washed into 20 μL sterile water in a 96 well PCR plate (Thermo Fisher Scientific, AB0600). After pipetting up and down 10 times, 5 μL cell suspension was spotted onto a new dropout agar plate (Nunc OmniTray). 15 μL of 40 mM sodium hydroxide was added into the PCR well with 15 μL residual cell suspension. The PCR plate was sealed and placed in a thermal cycler for boiling: 3 cycles of 98°C 3 min, 4°C 1 min. 50 μM primers were prefilled into a 384-well polypropylene plate (Labcyte, PP-0200). 0.2 μL of boiled yeast lysate and 30 nL of primer were used in a 5 μL GoTaq Green (Promega, M7123) PCR system. An acoustic liquid handler (Labcyte, Echo 550) was used for transferring primers and template into a 384 PCR plate (Greiner Bio-One, 785201).

#### Meiotic Recombination-mediated Assembly

Two semi-synthetic *synIV* strains with opposite mating types were mated on a YPD agar plate for 4-6 hours. Zygotes were pulled by using a tetrad dissection microscope (Singer Instruments, Sporeplay+). After two days of growth at 30°C, diploid cells were inoculated into GNA presporulation medium (5% dextrose, 3% Nutrient Broth, 1% yeast extract) for at least two doublings’ growth. Yeast cells were washed five times with sterile water, resuspended in 1x sporulation medium (1% w/v potassium acetate, 0.005% w/v zinc acetate dihydr*a*te), and subjected to sporulation for at least 5 days at 25°C. Tetrads were dissected on YPD plates; number of tetrads dissected was based on estimation of the single crossover frequency in the chromosome. Spores were cultured at 30°C on YPD for two days and replica-plated to drop out media to determine genotypes. Spores with desired genotype were further verified by PCRTagging to make sure the two synthetic chromosome segments were fully combined.

#### Transcript profiling

A single yeast colony was inoculated in 3 mL of YPD at 30°C with rotation overnight. Overnight yeast culture was diluted to A_600_=0.1 into 5 mL of fresh YPD liquid medium. Yeast cells were immediately harvested when A_600_ reached to 0.8-1.0. Total RNA was extracted using the RNeasy extraction kit (QIAGEN, 74106). 1 μg of total RNA was used as input for RNA library preparation (NEB E7770, E7490). RNA library was sequenced via Illumina 75 cycles high output kit on an Illumina NextSeq 550. Samples were first processed by removing the Illumina barcodes and adaptor sequences using Trimmomatic v0.33^63^. Processed reads were then aligned to the S288C transcriptome reference and a custom *synIV* transcriptome reference (in which chromosome *IV* transcripts had been replaced with their synthetic versions) using the program Kallisto v0.43.0^64^. All data analyses were carried out in Rstudio v1.1.456 (RStudio Team (2020). RStudio: Integrated Development for R. RStudio, PBC, Boston, MA URL http://www.rstudio.com/), using the Sleuth v0.30.0^65^ and ggplot2 v3.3.2 (Wickham H (2016). ggplot2: Elegant Graphics for Data Analysis. Springer-Verlag New York. ISBN 978-3-319-24277-4, https://ggplot2.tidyverse.org.) packages. For each sample we calculated log2 fold change values and tested for significance using Wald’s test and corrected for multiple testing with the false discovery rate adjusted p-value, using the Benjamini-Hochberg method.

#### Whole genome sequencing

Overnight cultured yeast cells were harvested by centrifugation at 3000xg for 2 min at room temperature. Yeast genomic DNA was extracted by using Fungi/Yeast Genomic DNA Isolation Kit (Norgen, 27300). The concentration of the genomic DNA was determined by Qubit dsDNA HS Assay Kit (ThermoFisher Scientific, Q32854). Approximately 300 ng of yeast genomic DNA was used as input for DNA library preparation. Whole genome sequencing libraries were prepared using the NEBNext Ultra II FS DNA library prep kit (NEB, E7805). Whole genome sequencing libraries were sequenced using an Ilumina NextSeq 550 75 cycles high output kit.

#### Genome sequencing analysis

The genomic sequence of synIV was analyzed using the Synthetic Yeast sequencing (SYseq) pipeline (Stracquadanio, G. et al, in preparation). Briefly, Illumina paired-end reads were first preprocessed to trim adapters and remove low quality bases, and then aligned to the BY4741 reference genome were the wild-type chromosome IV was replaced by the synthetic counterpart. The resulting alignments were used to identify single nucleotide variants (SNVs) and short indels using a new freebayes protocol^66^, whereas structural variants and copy number changes were identified by combining GRIDSS^67^ with a new copy number calling algorithm specifically designed to work on haploid strains. Finally, mutational and coverage data were organized into VCF and bigWig files for downstream analysis and easy visualization.

#### Hi-C library preparation and data analysis

Yeast cells were inoculated into 10 mL of YPD overnight, and the next day 10^9^ cells (approximately 80-100 OD) were subcultured into 150 mL YPD for 2.5 hours of growth at 30°C. Cells were crosslinked by 3% [v/v] formaldehyde for 20 min at room temperature and subsequently quenched with glycine for 15 min at 4°C. Crosslinked cells were harvested and suspended in 10 mL spheroplast buffer (1M sorbitol, 5 mM DTT, 250 U zymolyase 100T (US Biological, Z1004)) for 40 min incubation at 30°C. Spheroplasts were washed with 10 mL of 1M sorbitol once and resuspended in 2 mL of 0.5% SDS at 65°C for 20 min. *Mbo*I (NEB, R0147) was used for overnight digestion of all the genomic DNA from fixed yeast cells at 37°C. The digestion product was centrifuged at 18000xg for 20 min and the pellet was subsequently suspended in 200 μL cold water. DNA sticky ends were filled with biotin-14-dCTP (Invitrogen, 19518018) by Klenow enzyme (NEB, M0210L) at 37°C for 80 min. Biotinylated DNA was ligated with T4 ligase (Thermo Scientific, EL0014) at room temperature for 2 hours. Ligation product was reverse cross-linked by 0.5 mg/mL proteinase K (Thermo Scientific, EO0492) in 25 mM EDTA buffer at 65°C for 4 hours. The reverse crosslinked sample was purified using the large fragment DNA recovery kit (Zymo Research, D4046). Purified Hi-C library was used as input material for Illumina sequencing library prep kit (NEB, E7805). DNA library was sequenced using an Illumina NextSeq 550 150-cycle high output kit. Hi-C data were analyzed using a multistep, parameter-free approach as previously described^68,69^. Briefly, Hi-C reads were mapped to the reference with Bowtie2. Normalized contact maps were 5 kb-binned, and were plotted using MatLab2018. PyMol was used for the 3D projection.

#### Immunoblotting

Middle-log-phase yeast cells were harvested by centrifugation at 3000xg for 3 min at 4°C. Yeast cells were suspended in 200 μL of lysis buffer (20 mM HEPES pH7.4, 0.1% Tween20, 2 mM MgCl_2_, 300 mM NaCl, 1× protease inhibitor (Roche)). Approximately 100 μL of 0.5 mm glass beads were added into the cell suspension. Tubes were subjected to a Retsch MM 400 mixer 3 times with 1 min shaking. Centrifuge the tubes at 12000xg for 10 min at 4°C. 30 μL of supernatant was transferred to a new tube, and mixed with 10 μL of 4 LDS loading buffer (Invitrogen, NP0007). Samples were heated to 70°C for 10 min. 10 μL samples were loaded onto an SDS-PAGE gel for immunoblotting. Nop1 antibody (Invitrogen, MA110025), and histone H3 antibody (Abcam, 1791) were used for blotting the target proteins. Goat anti-mouse (LI-COR Biosciences, 926-32210) and goat anti-rabbit (LI-COR Biosciences, 926-68071) secondary antibodies were used for blotting Nop1 and H3 primary antibodies respectively. A LI-COR Odyssey instrument was used to develop the blot images^70^.

#### RT-qPCR

Approximately 2×10^7^ mid-log-phase yeast cells were harvested by centrifugation at 3000xg for 3 min. Total RNA was extracted following the instructions of QIAGEN RNeasy kit (QIAGEN, 74106). 1 μg of total RNA was used for the first-strand cDNA synthesis by SuperScript III kit (Invitrogen, 18080051). LightCycler 480 SYBR Green I Master Mix (Roche, 04887352001) was used for the qPCR reaction. LightCycler 480 instrument was used to perform the qPCR.

#### Pulsed-field gel electrophoresis

A single yeast colony was inoculated in 10 mL of YPD medium and cultured for 2 days at 30°C. Approximately 30 mg of yeast cells were harvested by centrifugation. 12 μL of zymolyase solution (25 mg/mL zymolyase 20T [US Biological, Z1000] in 10 mM KH_2_PO_4_ pH 7.5) and 270 μL of melted low melting point agarose were added to the cell pellet. The cell suspensions were mixed by pipetting, and yeast cell plugs were made using BioRad plug molds. After cooling for 1 hour at room temperature, solidified plugs were transfer into 15 mL tubes containing 1 mL of digestion buffer (0.5 M EDTA, 10 mM Tris, pH 7.5). Yeast cell plug containing tubes were gently inverted to mix and incubated at 37°C overnight. The next day, 400 μL proteinase K solution (5% sarcosyl, 5 mg/mL proteinase K in 0.5 M EDTA, pH 7.5) was added to the tube for 5 hours of incubation at 50°C. Plugs were washed in TE pH8.0 three times. Plugs were loaded onto pulsed field electrophoresis system using 1% agarose gels run in 0.5xTBE buffer (BioRad, CHEF Mapper XA) for chromosome separation.

#### Swapping TAG stop codon to TAA by SpCas9-NG in yeast

The SpCas9-NG coding sequence was amplified from the pX330-SpCas9-NG plasmid (Addgene, 117919). The yeast *TEF1* promoter and *CYC1* terminator were cloned to assemble an SpCas9-NG transcription unit in pRS413 backbone (with *HIS3* marker), creating plasmid pWZ399. Guide RNA (gRNA) was designed by selecting the 19 bp upstream of TAG stop codon plus a thymine (T) as the 20^th^ nucleotide. Guide RNA was cloned into a 2-micron plasmid (pRS426, with *URA3* marker) under *SNR52* promoter as described^71^. The SpCas9-NG transcription unit plasmid was transformed into yeast to pre-establish SpCas9-NG in the cells. 50 ng of gRNA plasmid and 200 ng donor DNA fragment were co-transformed into the SpCas9-NG containing yeast strain to enable G to A editing. Transformants were grown on SC–Ura–His plate. G to A editing was verified by Sanger sequencing.

#### Chromosome stability assay

Haploid *synIV* and wild type strains were used for *URA3* gene insertion to the left arm and *KanMX4* gene insertion to the right arm. Heterozygous diploid strains were created by mating on YPD at 30°C for 6 hours followed by zygotes isolation using a tetrad dissection microscope. Heterozygous diploid strains with *URA3* and *KanMX4* either on synthetic or wild-type chromosome were grown in 5 mL YPD culture at 30°C until the saturation stage at which time 1:1000 dilution was made to a fresh 5 mL YPD culture. After approximately 100 generations, yeast cells were plated on YPD and 5-FOA plates for colony counting. Colonies on 5-FOA were replica plated to G418 for loss of heterozygosity evaluation.

#### Zinc Finger binding affinity assay

A bacterial one-hybrid assay^41^ was used to evaluate the binding activity between synthetically designed zinc finger proteins and the loxPsym sequence. Briefly, zinc finger candidates were fused with the omega subunit of *E. coli* RNA polymerase. The 34 bp loxPsym sequence was cloned upstream of *HIS3* on a reporter plasmid. *E. coli* cells with omega subunit knockout harboring the aforementioned two plasmids were plated on 2xYT medium or medium lacking histidine and supplemented with different concentrations of 3-AT.

#### Synthetic chromosomes tethering

BY4741*, synIV, synIII* strains transformed with HEH1/HEH2-only, HEH1-ZF/HEH2-ZF, or ZF-only plasmids were grown in SC–Leu–Ura medium. For chromosome tethering assays on solid medium, yeast cells in early log phase were spotted on SC–Leu–Ura medium supplemented with 10 nM estradiol. For chromosome tethering assays in liquid medium, yeast cells were diluted in 96 well culture plate with starting A_600_ about 0.2. A_600_ was measured using a plate reader (Cytation 5, BioTek) for every 10 min, with double-orbital continuous shaking settings.

## Supplementary information

**Fig. S1.**
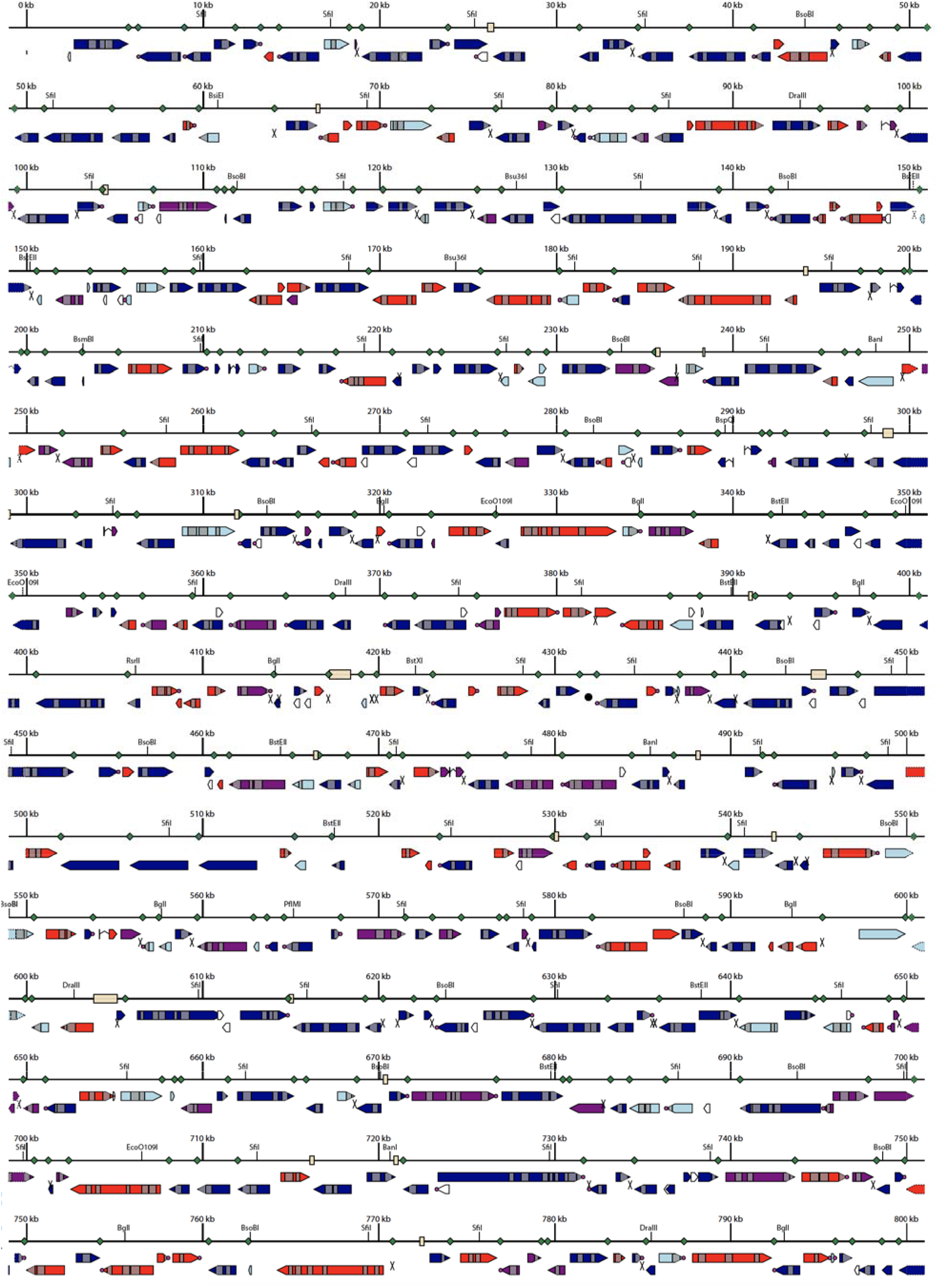
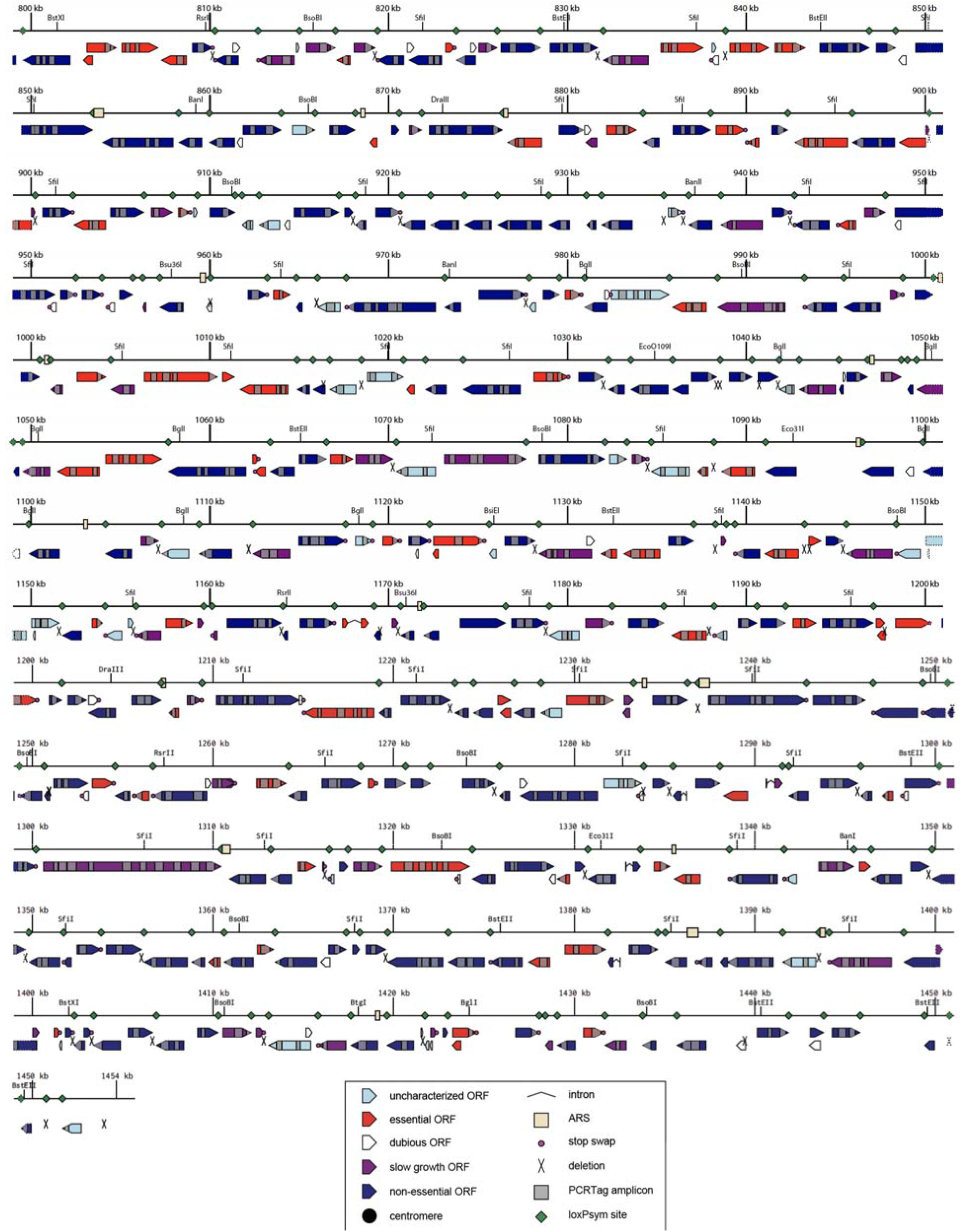
Schematic illustration of all design features across the 1454 kb *synIV*. Related to Fig. 1. Features are shown in the bottom box.

**Fig. S2.**
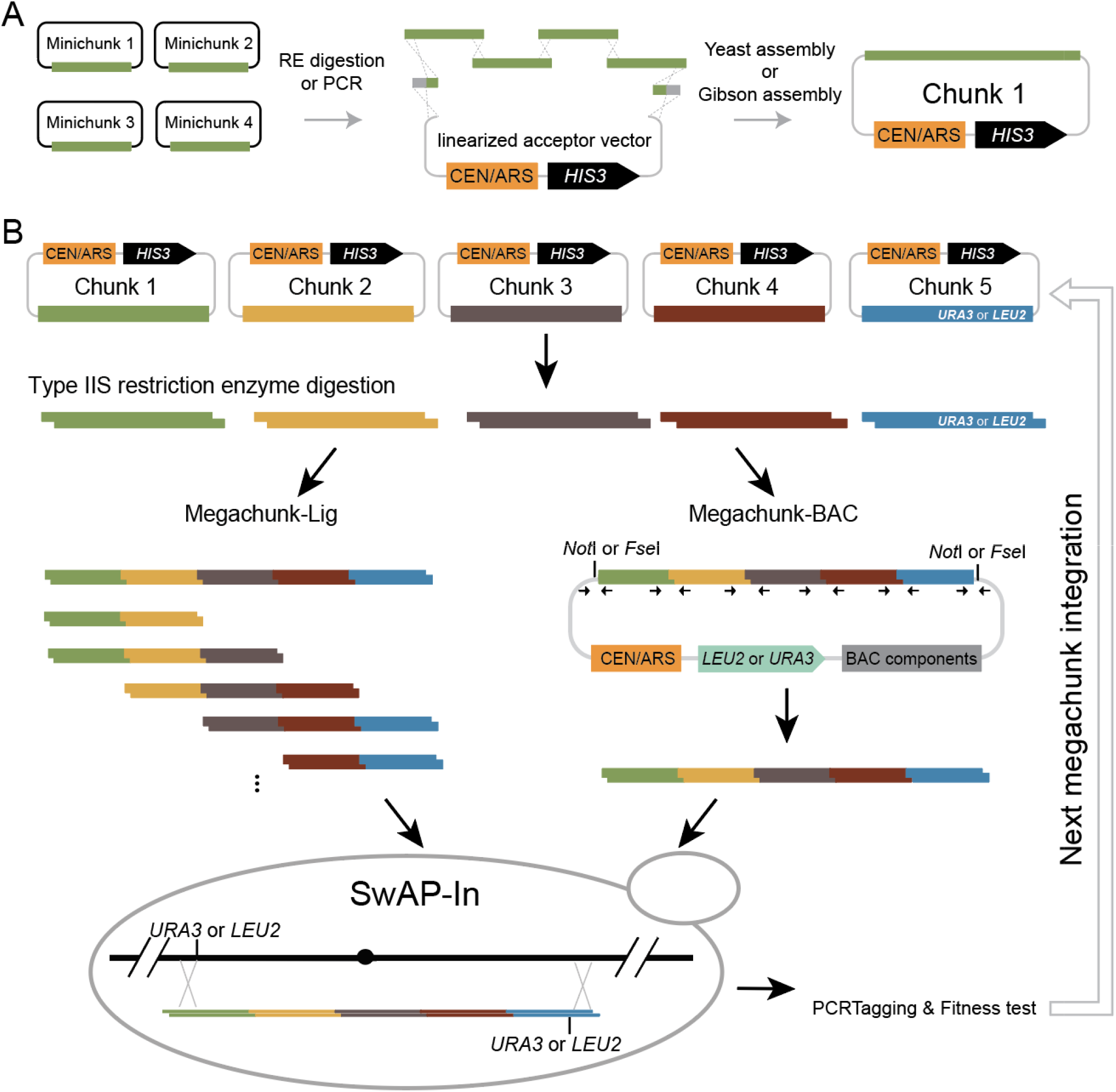
Schematic illustration of chunk, megachunk cloning and megachunk integration. Related to. Fig. 1. (**A**) Minichunk DNAs were assembled into chunk DNA via yeast homologous recombination or Gibson assembly. (**B**) Megachunk integration using conventional megachunk ligation method and new megachunk assembly method. The chunk DNAs are shown by lines with different colors. Megachunks were assembled into pWZ212 (*URA3*) or pWZ237 (*LEU2*). BAC, bacterial artificial chromosome.

**Fig. S3.**
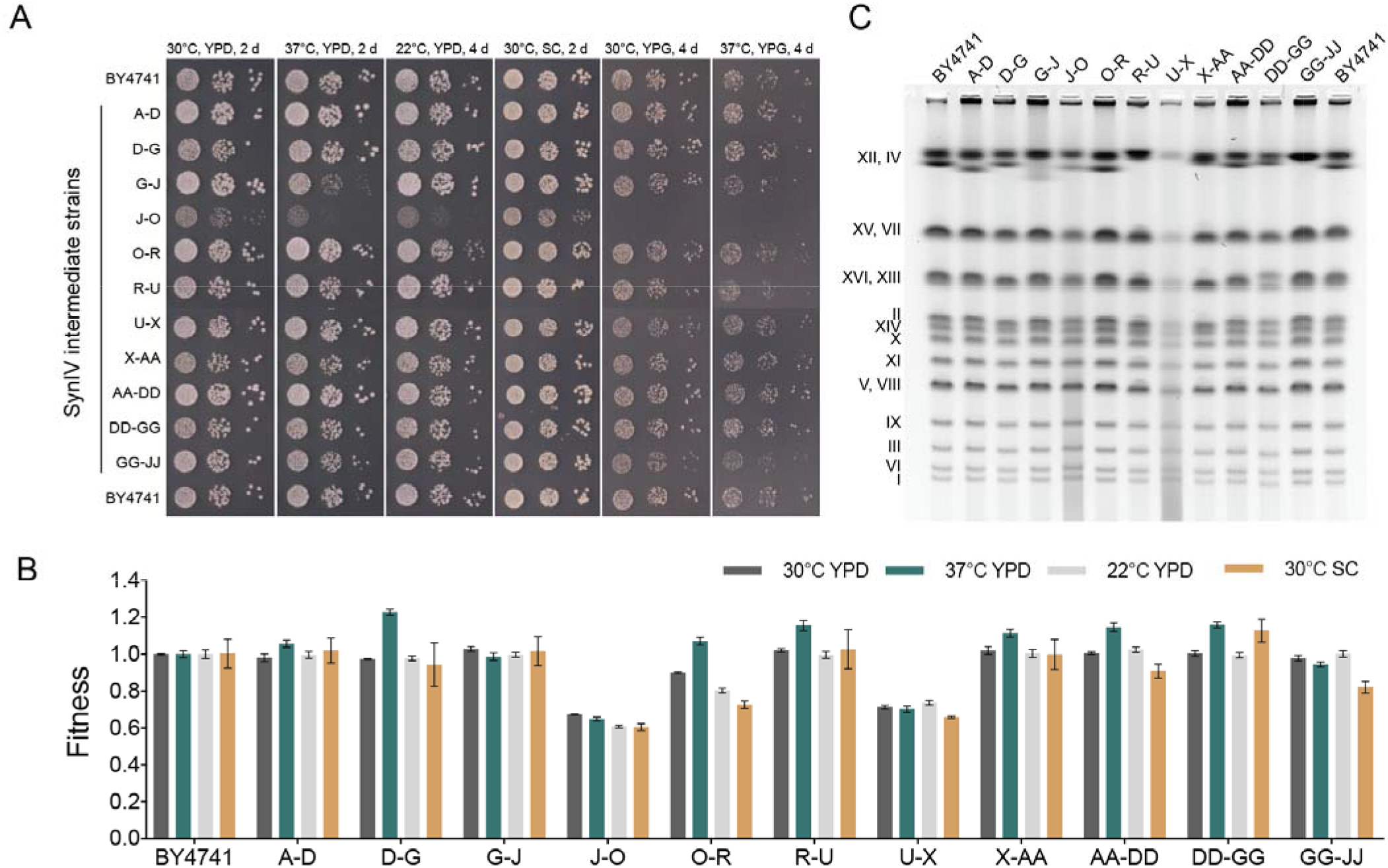
*SynIV* intermediate strains fitness and karyotyping. Related to Fig. 1. (**A**) Ten-fold serial dilution spot assay analysis of 11 semi-*synIV* strains under six different solid media growth conditions. YPD, yeast extract, peptone, dextrose. YPG, yeast extract, glycerol. SC, synthe ic complete. (**B**) Growth fitness of 11 semi-*synIV* strains measured in liquid medium culture condition. (**C**) Karyotyping analysis of 11 semi-*synIV* strains by pulse-field gel electrophoresis. Bio-Rad CHEF mapper auto algorithm was used, low molecular weight, 200 kb; high molecular weight, 1500 kb; voltage, 6 V/cm; total electrophoresis time was 24 h.

**Fig. S4.**
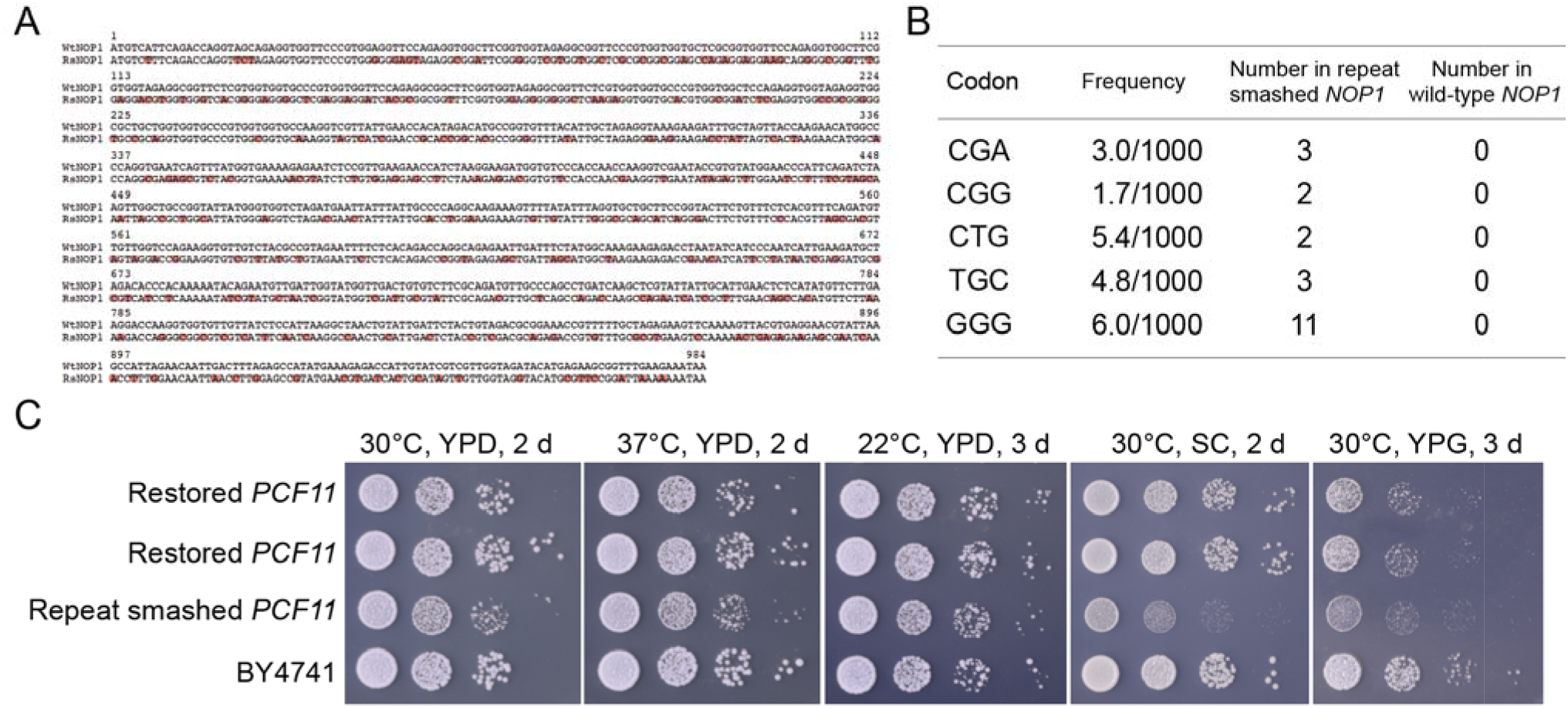
*NOP1* debugging. Related to Fig. 2. (**A**) Sequence alignment of wild-type and repeat-smashed *NOP1*. (**B**) Number of non-optimal codons present in wild-type and repeat-smashed *NOP1*. (**C**) Serial dilution assay showing growth defects when strain carries repeat-smashed *PCF11*.

**Fig. S5.**
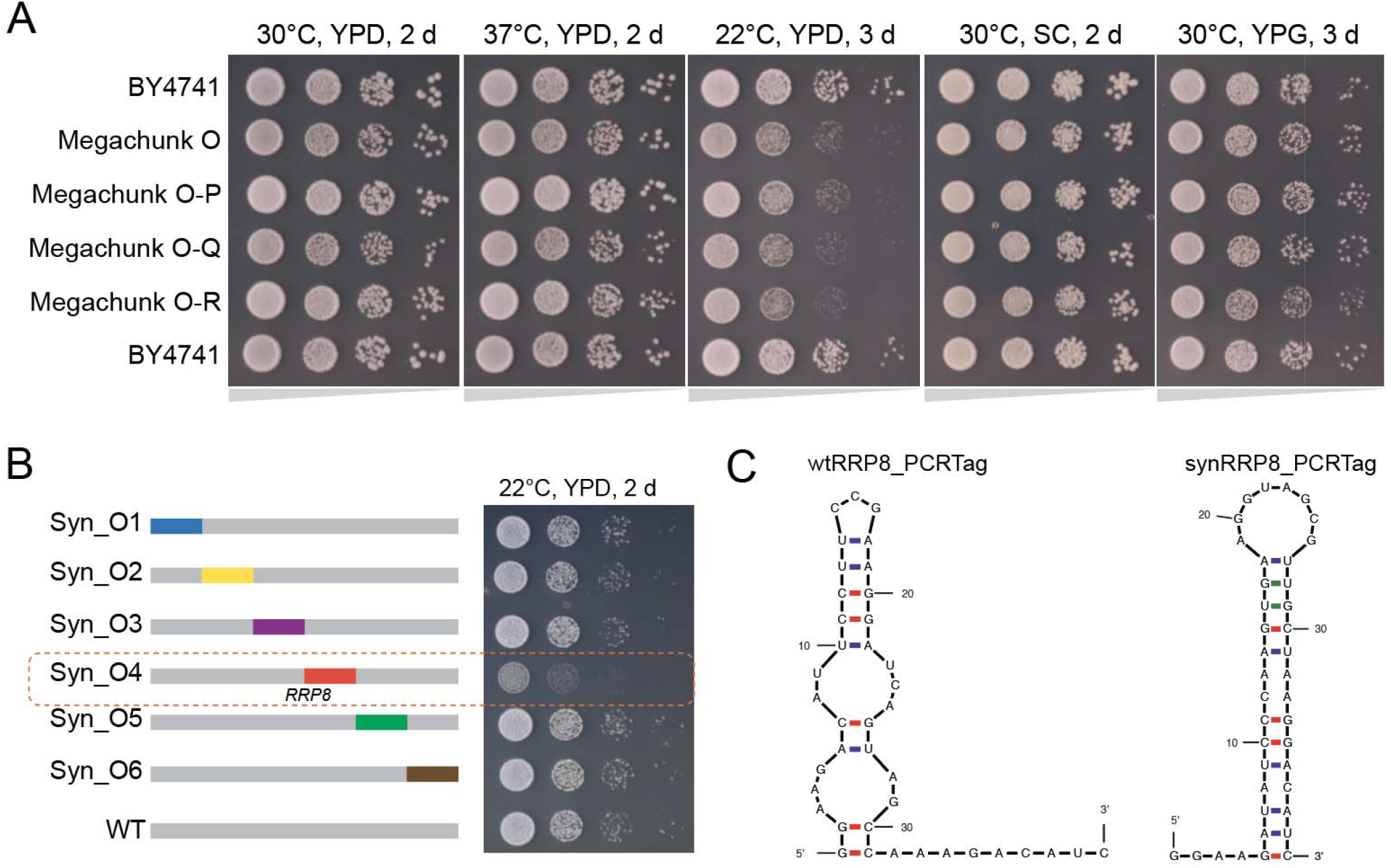
*RRP8* debugging. Related to Fig 2. (**A**) Growth fitness test of a series of megachunk strains. (**B**) Strategy used to map the bug in megachunk O. (**C**) mRNA secondary structure prediction by mFold. Wild-type or synthetic *RRP8* PCRTag plus 6 bp flanking bases were used for the prediction. dG is −9.67 for wtRR8_PCRTag, dG is −8.93 for synRR8_PCRTag.

**Fig. S6.**
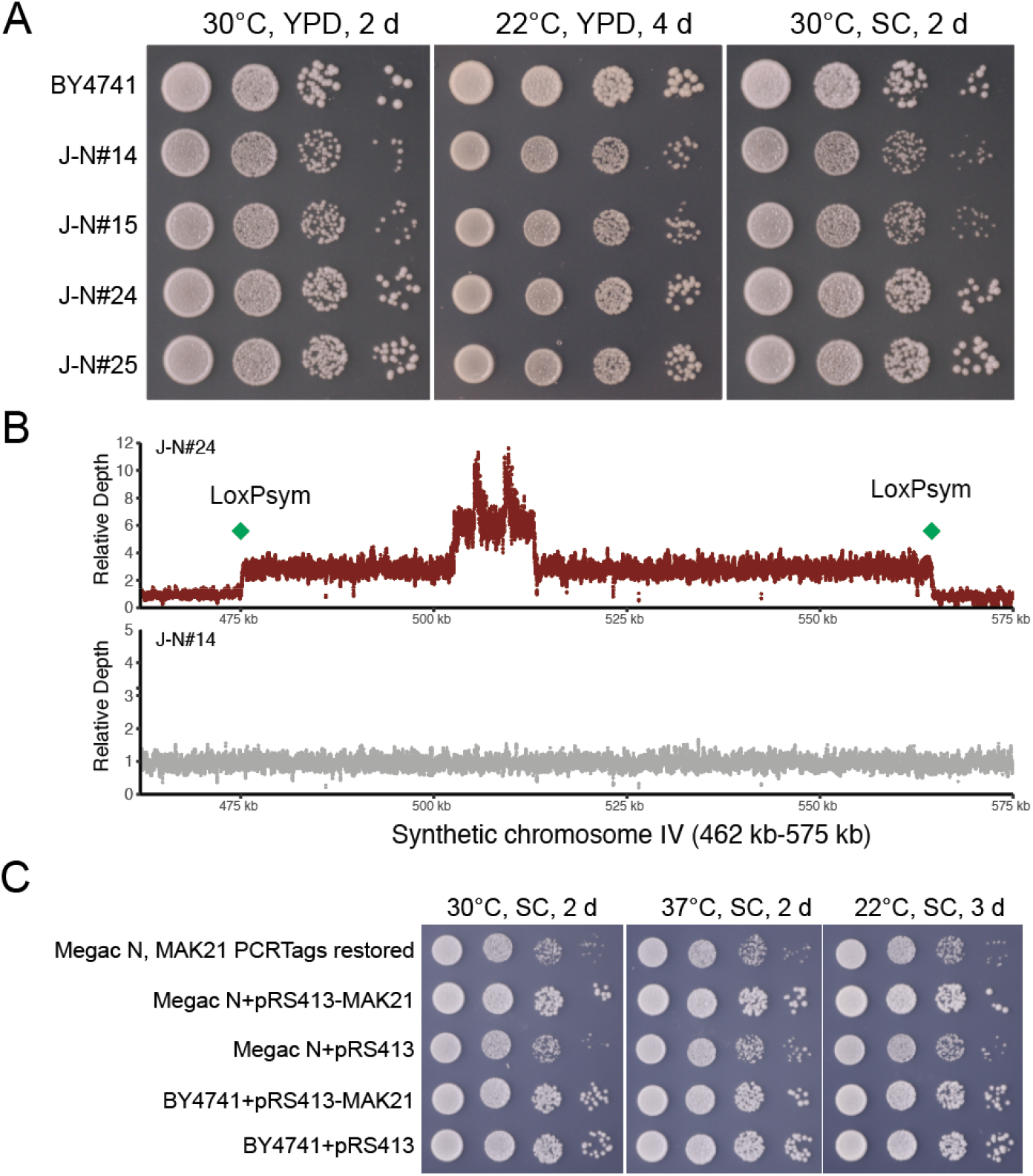
*MAK21* debugging. Related to Fig. 2. (**A**) Serial dilution assay showing heterogenous growth fitness. (**B**) Sequencing read depth plot of megachunk J integrants with or without intrachromosomal duplication. (**C**) Growth fitness of megachunk N strain was restored by a plasmid harboring wild-type *MAK21* gene (500 bp upstream for promoter and 200 bp downstream for terminator), rather than by restoration of the synthetic PCRTags of *MAK21*.

**Fig. S7.**
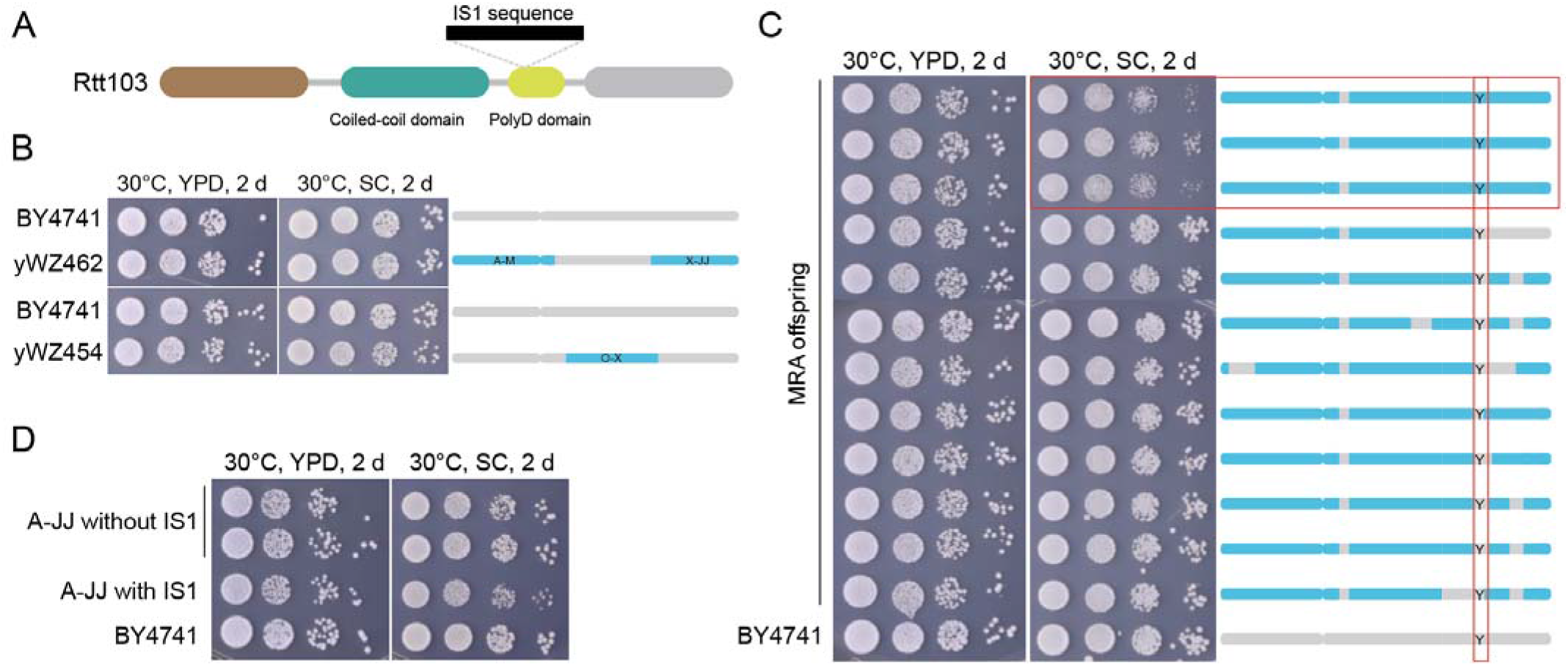
*RTT103* debugging. Related to Fig.2. (**A**) A 777-bp *E. coli* IS1 transposon was inserted in the poly aspartic acid domain of *RTT103*, disrupting the gene. (**B**) Serial dilution dot assay showing neither yWZ462 (contains megachunk A-M and X-JJ) nor yWZ454 (contains megachunk O-X) showed growth defect on SC medium. (**C**) Severe growth defect was observed among some of the MRA offspring. Genotyping PCR identified the synthetic “pattern” in each offspring. Megachunk Y containing strains consistently showed growth defects on SC medium. (**D**) Serial dilution assay shows fitness restoration after removing IS1 sequence.

**Fig. S8.**
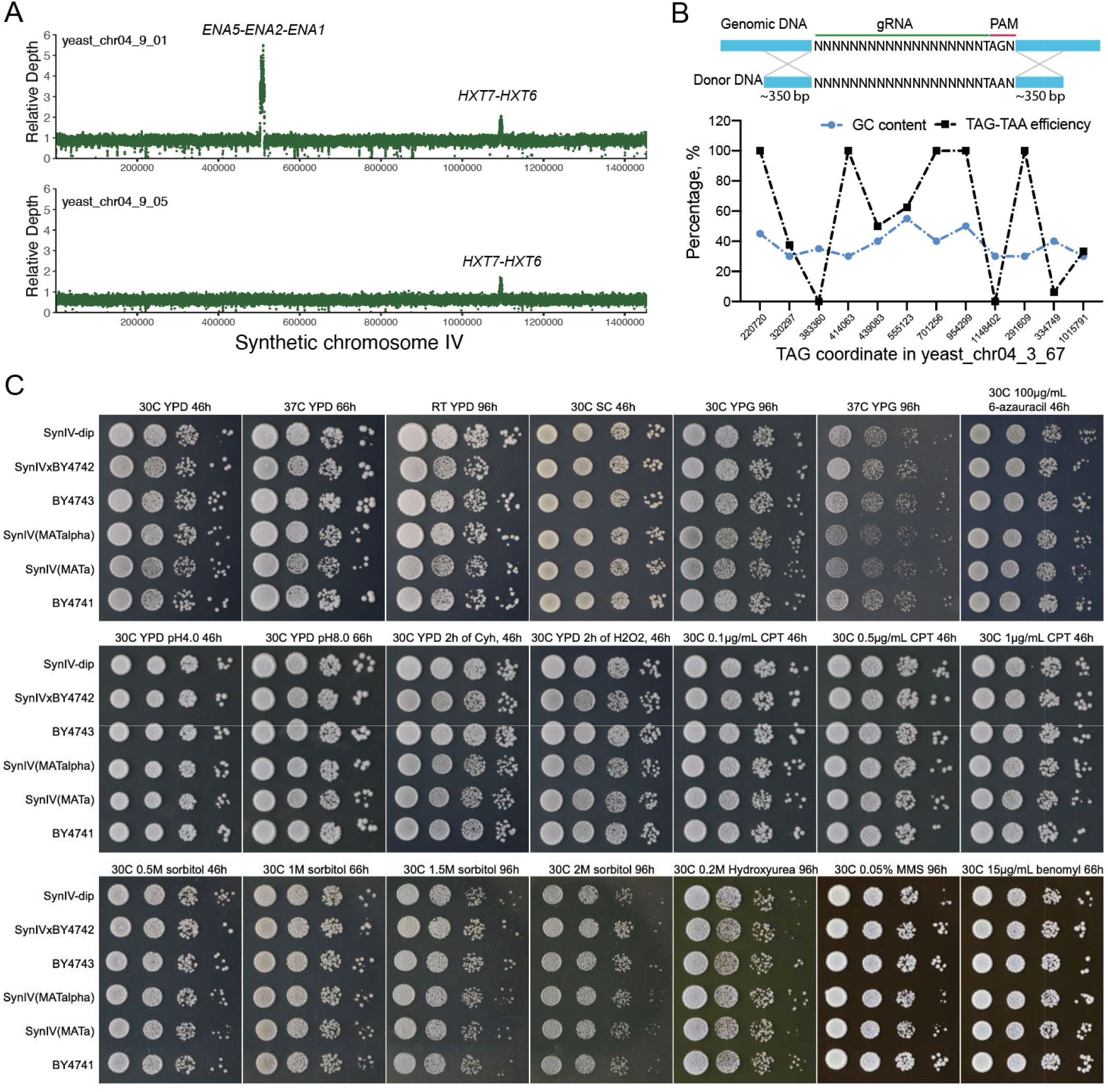
*SynIV* sequence correction and growth fitness tests. Related to Fig. 3. (**A**) Whole genome sequencing read depth plot of two *synIV* living strains. X axis, *synIV* coordinate, y axis, relative read depth. The small peak at pos. ∼1,100,000 is *HXT7-HXT6* duplication. (**B**) SpCas9-NG enabled TAG to TAA editing. SpCas9-NG enabled TAG-TAA editing efficiency was computed by the number of Sanger sequencing verified clones divided by the total sequenced clones. (**C**) Haploid, heterozygous diploid and homozygous diploid *synIV* growth under various stress conditions.

**Fig. S9.**
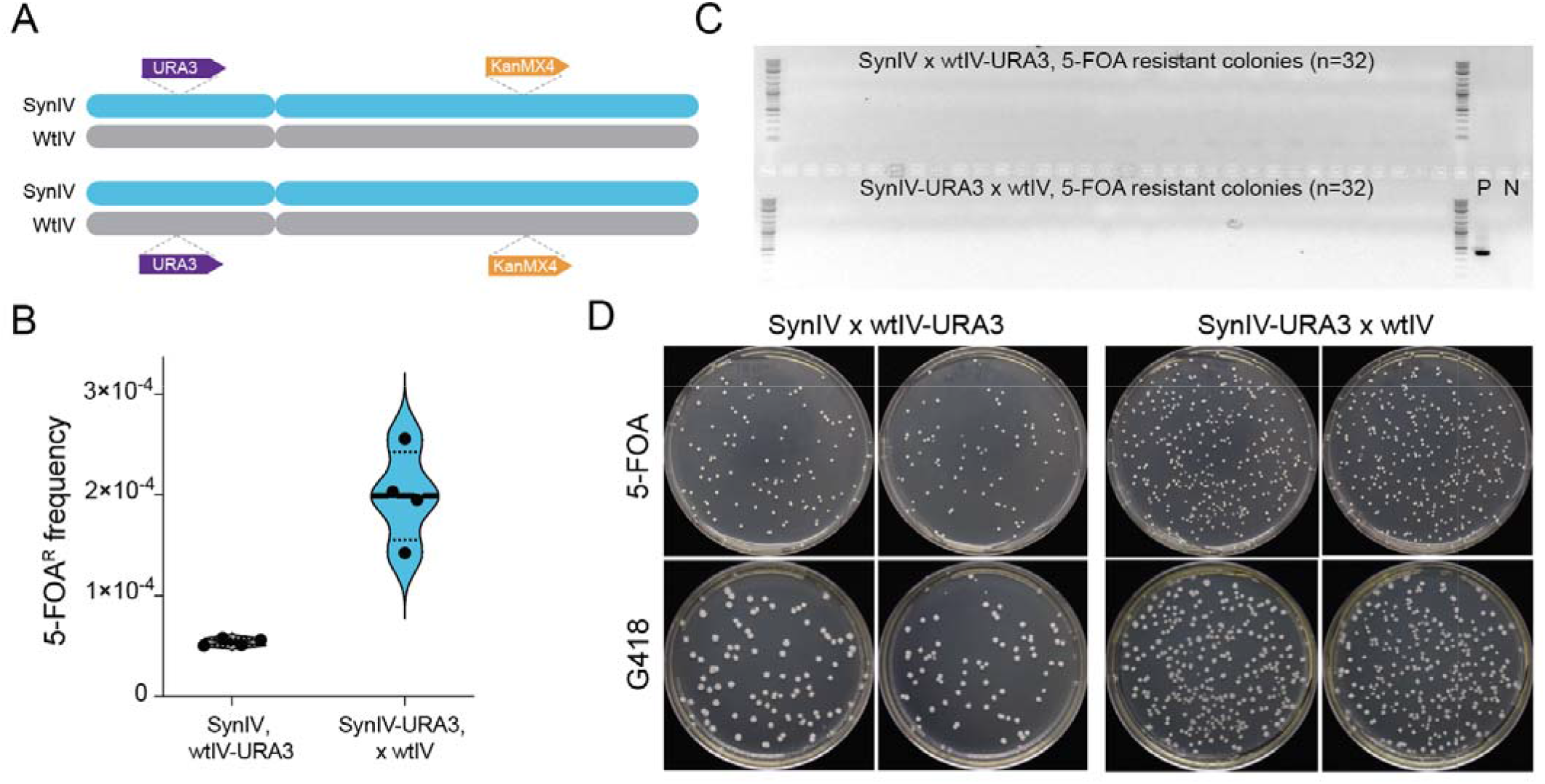
*SynIV* stability evaluation. Related to Fig. 3. (**A**) Schematic diagram showing the *URA3* and *KanMX4* insertions in the heterozygous diploid *synIV* strains. (**B**) 5-FOA resistance frequency comparison between the two heterozygous diploid *synIV* strains in A. (**C**) *URA3* genotyping PCR analysis for the 5-FOA resistant colonies. (**D**) Representative images for colonies on 5-FOA plates and their G418 replica plates.

**Fig. S10.**
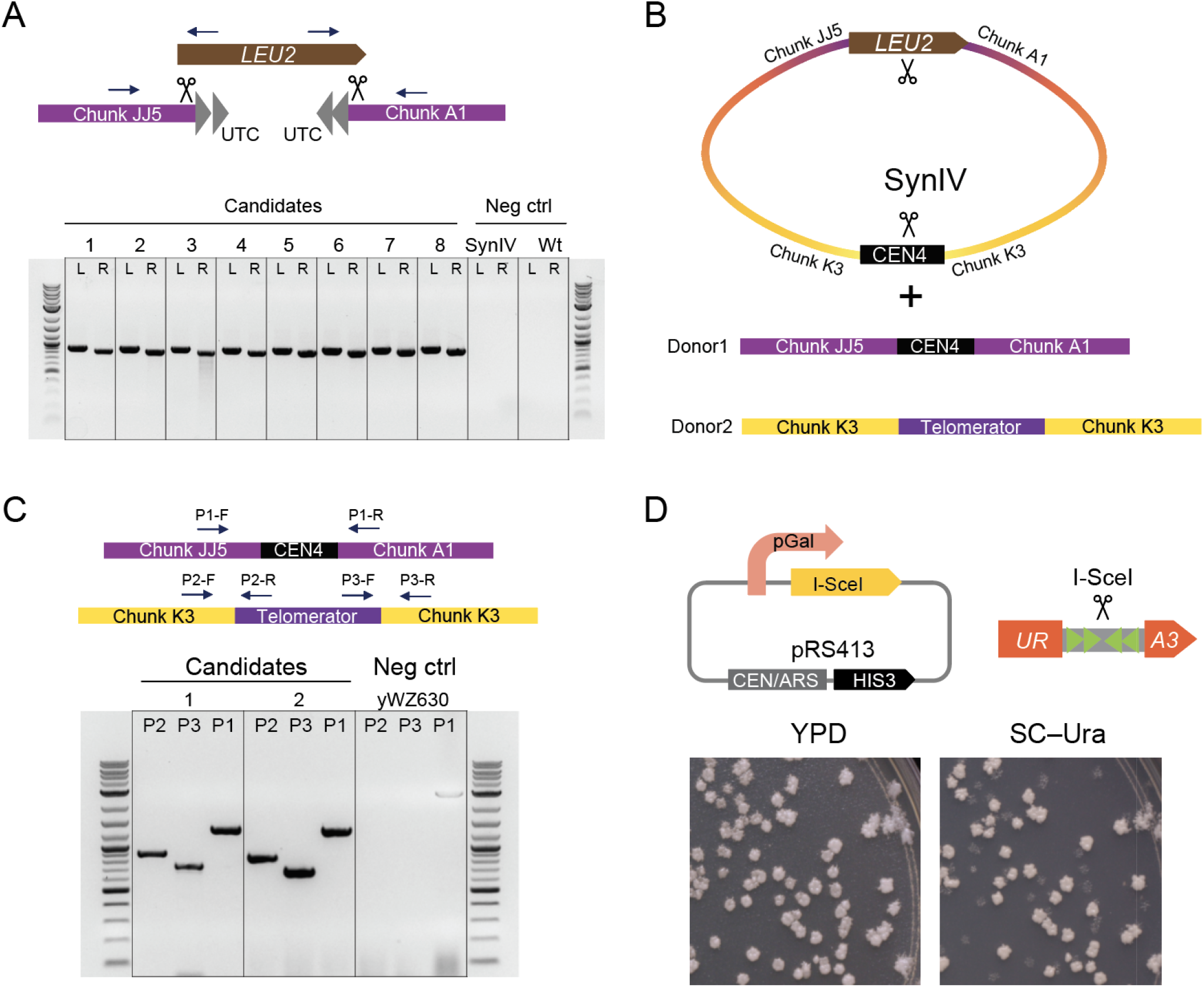
Detailed engineering of “inside-out” *synIV*. Related to Fig. 4. (**A**) *SynIV* circularization. Two telomeres were removed by CRISPR-Cas9-induced DNA double strand breaks. A *LEU2* gene with homology arms to chunk JJ5 on the left side and chunk A1 in the right side was provided as the donor template. Arrows indicate the junction PCR primer locations and directions., left junction; R, right junction. (**B**) Intra-chromosomal centromere relocation. Two DNA double strand breaks were introduced on *LEU2* gene and *CEN4* respectively, two repair donors (donor 1 and donor 2, DNA content was indicated in the figure) were co-transformed into the cells to facilitate the *CEN4* relocation. (**C**) Junction PCR verification of *CEN4* relocated synIV strains. (**D**) Inside-out *synIV* linearization by transiently expressing I-*Sce*I endonuclease.

**Fig. S11.**
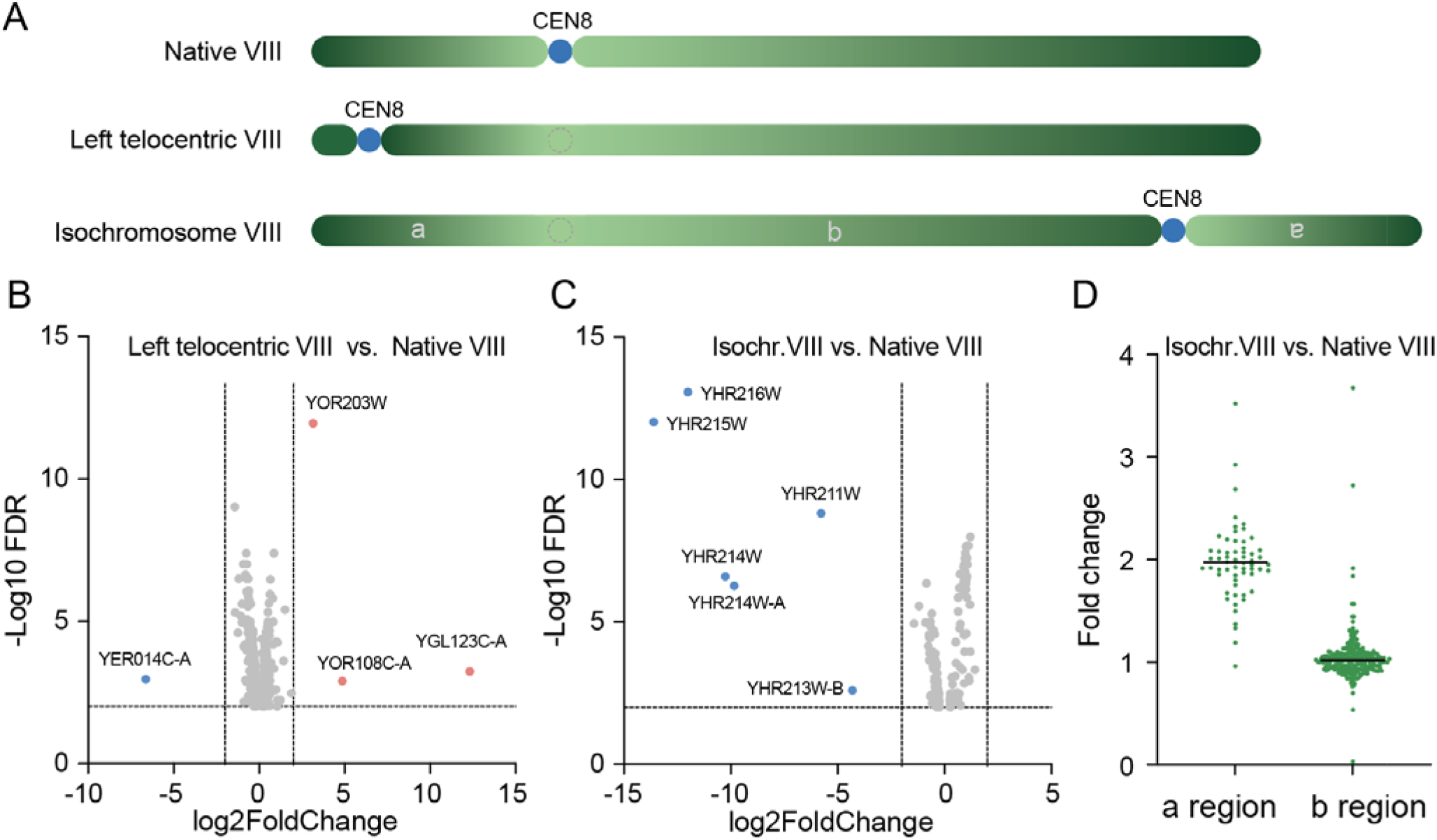
*CEN8* relocation did not alter chromosome VIII gene expression. Related to Fig. 4. (**A**) Schematic showing the three *CEN8* relocation strains. Gradient green bars indicate directionality of chromosome arms. Blue circles are *CEN8* core sequence, dotted circles showing native *CEN8* location in the isochromosome. (**B**) Volcano plot of differentially expression genes from left telocentric VIII vs. native VIII. (**C**) Volcano plot of differentially expression genes from isochromosome VIII vs. native VIII. Fold change cutoff is 4, FDR cutoff is 0.01. Note that ORFs YHR211-216 were lost in the formation of the isochromosome. (**D**) Left chromosome arm genes were upregulated by 2-fold in the isochromosome VIII due to copy number increase. Fold change cutoff is 4, FDR cutoff is 0.01.

**Fig. S12.**
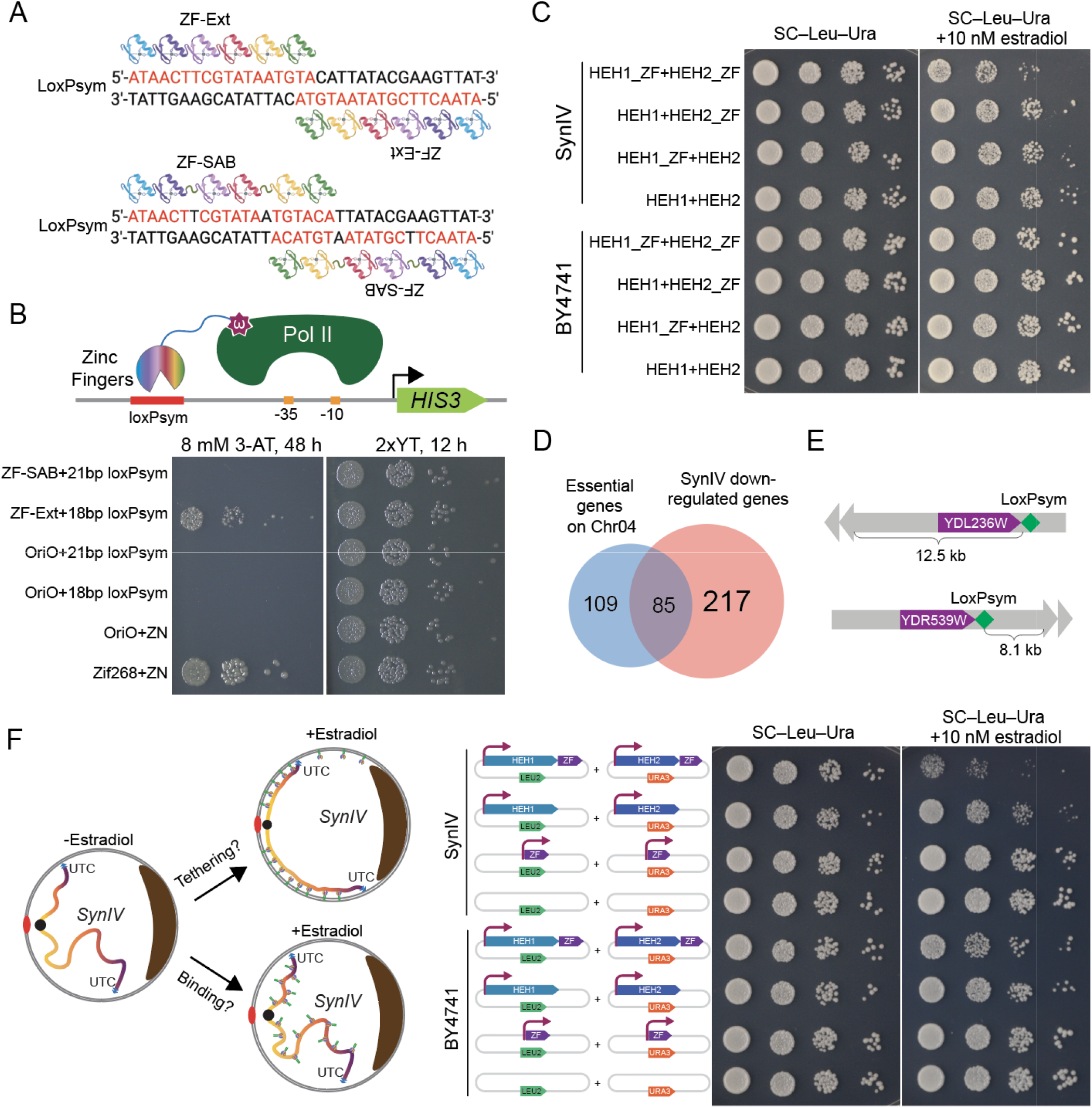
LoxPsym binding zinc finger design and *synIV* tethering. Related to Fig. 5. (**A**) Two synthetically designed zinc finger proteins for loxPsym site. ZF-Ext has six fingers binding to a consecutive 18 bp sequence on loxPsym, ZF-SAB has six fingers that skip one base pair every two fingers. Icons were created with BioRender.com. (**B**) Reporter assay testing the binding activity of synthetic zinc fingers to the loxPsym site. *E. coli* cells were co-transformed with *HIS3* reporter plasmid and zinc finger expression plasmid. *E. coli* transformants were spotted on 2xYT medium and histidine dropout medium supplemented with 8 mM 3-AT. (**C**) Ten-fold serial dilution spot assay for *synIV* and BY4741 containing indicated plasmids. Yeast cells were spotted on either SC–Leu-Ura medium or SC–Leu-Ura+10 nM estradiol. (**D**) Venn diagram showing chromosome IV essential gene list overlapping with downregulated *synIV* genes. (**E**) Distances between the universal telomere cap and the first downregulated gene’s loxPsym site on each chromosome arm. (**F**) Ruling out *synIV* repression is caused by zinc finger protein binding on the chromosome. Ten-fold serial dilution spot assay showing expressing ZF alone has no impact on strains’ growth.

**Fig. S13.**
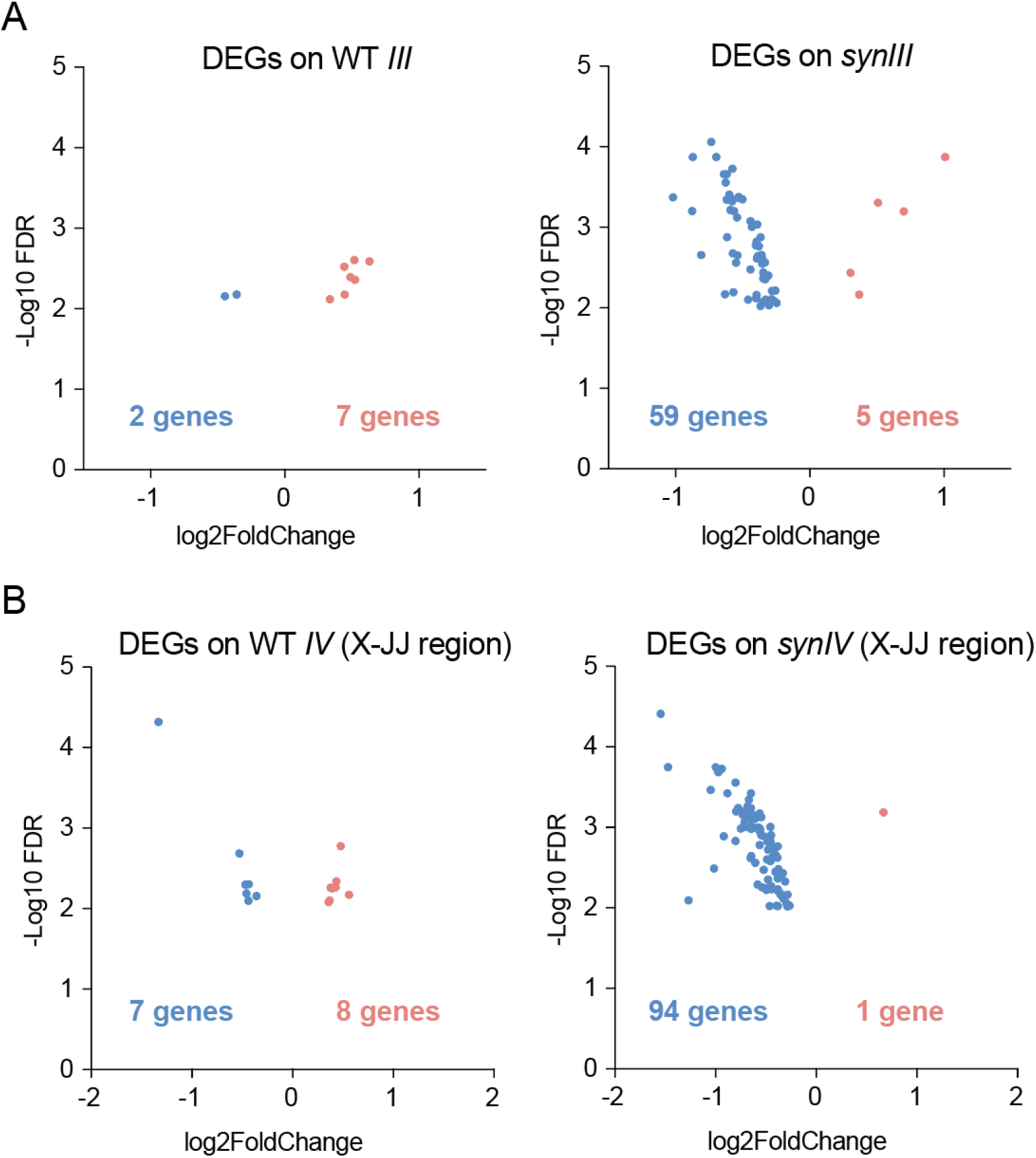
*SynIII* and *synIV X-JJ* tethering. Related to Fig. 6. (**A**) Volcano plots of differentially expressed *synIII* genes from HEH1-ZF/HEH2-ZF vs. HEH1/HEH2-only in WT (left panel) or *synIII* (right panel) strain. Fold change cutoff is 4, FDR cutoff is 0.01. (**B**) Volcano plots of differentially expressed *semi-synIV* megachunk X-JJ region genes (*YDR255C* to *YDR541C*) from HEH1-ZF/HEH2-ZF vs. HEH1/HEH2-only in WT (left panel) or *synIII* (right panel) strain. Fold chan e cutoff is 4, FDR cutoff is 0.01.

**Table S1.** Repeat smashed gene list. Related to Figure 1A.

| Essential genes | Nonessential genes |
| --- | --- |
| *YDL014W (NOP1)* | *YDR077W (SED1)* |
| *YDR420W (HKR1)* | *YDR534C (FIT1)* |
| *YDR228C (PCF11)* | *YDR150W (NUM1)* |
|  | *YDL037C (BSC1)* |

**Table S2.**
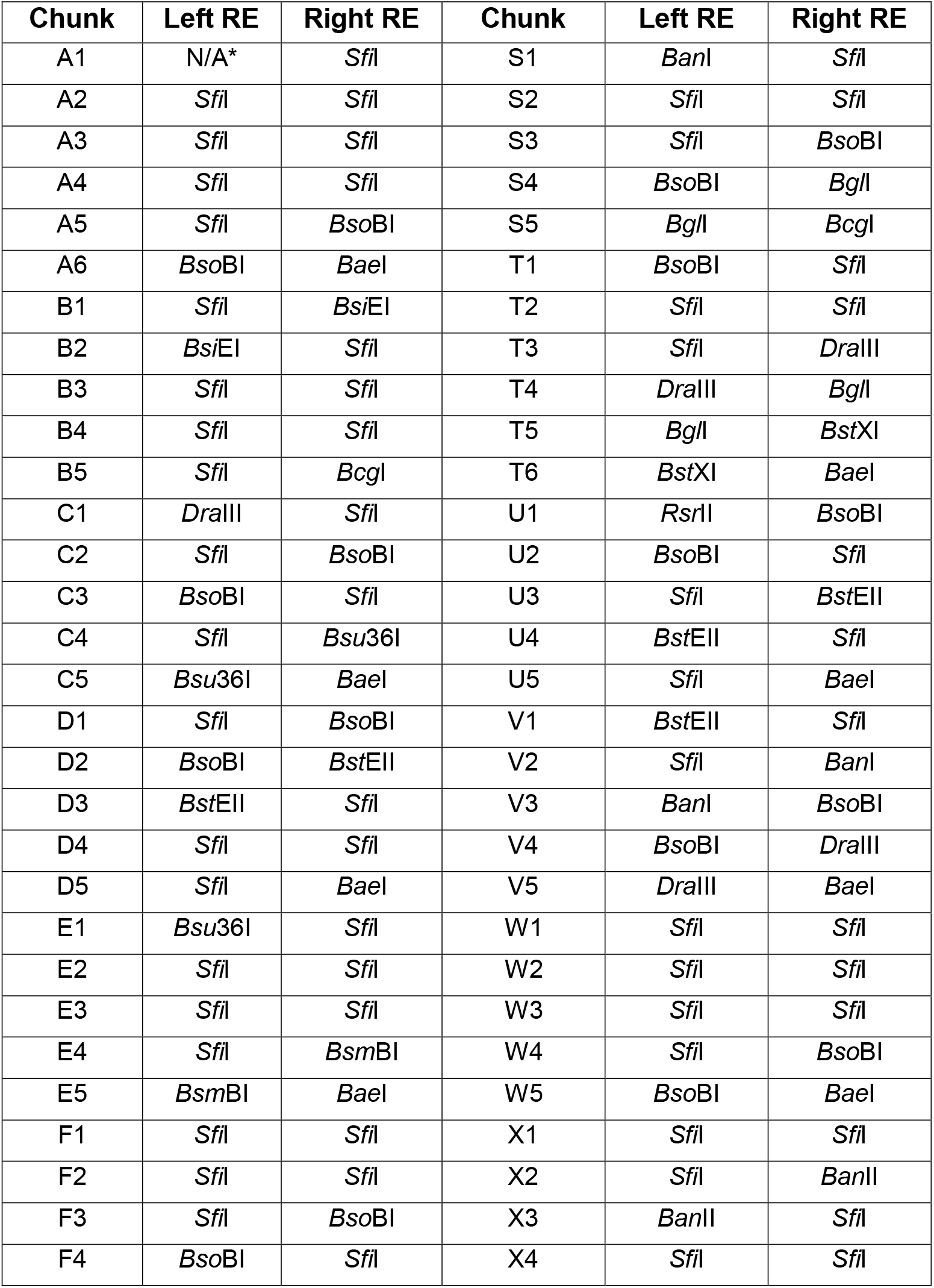
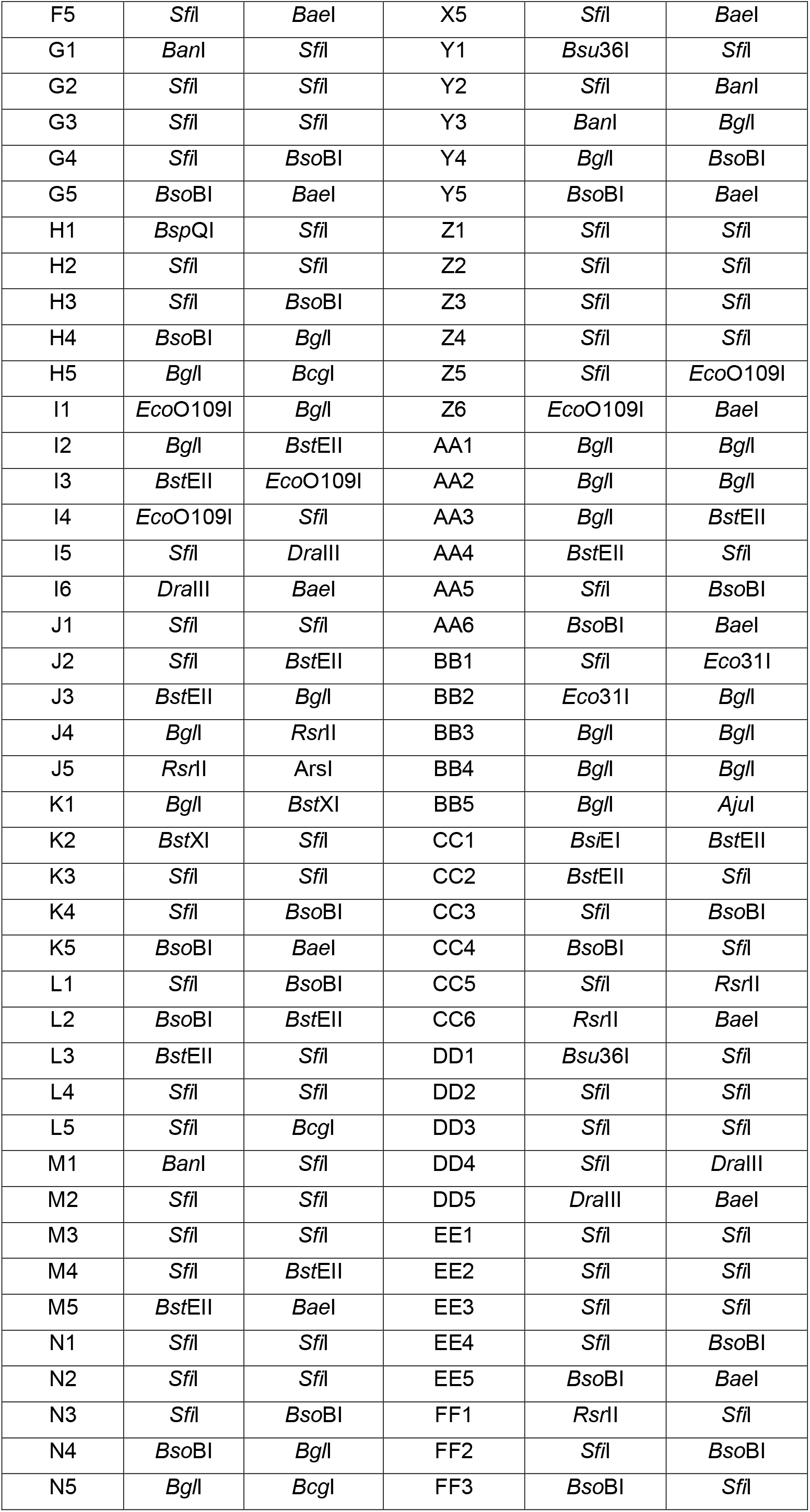
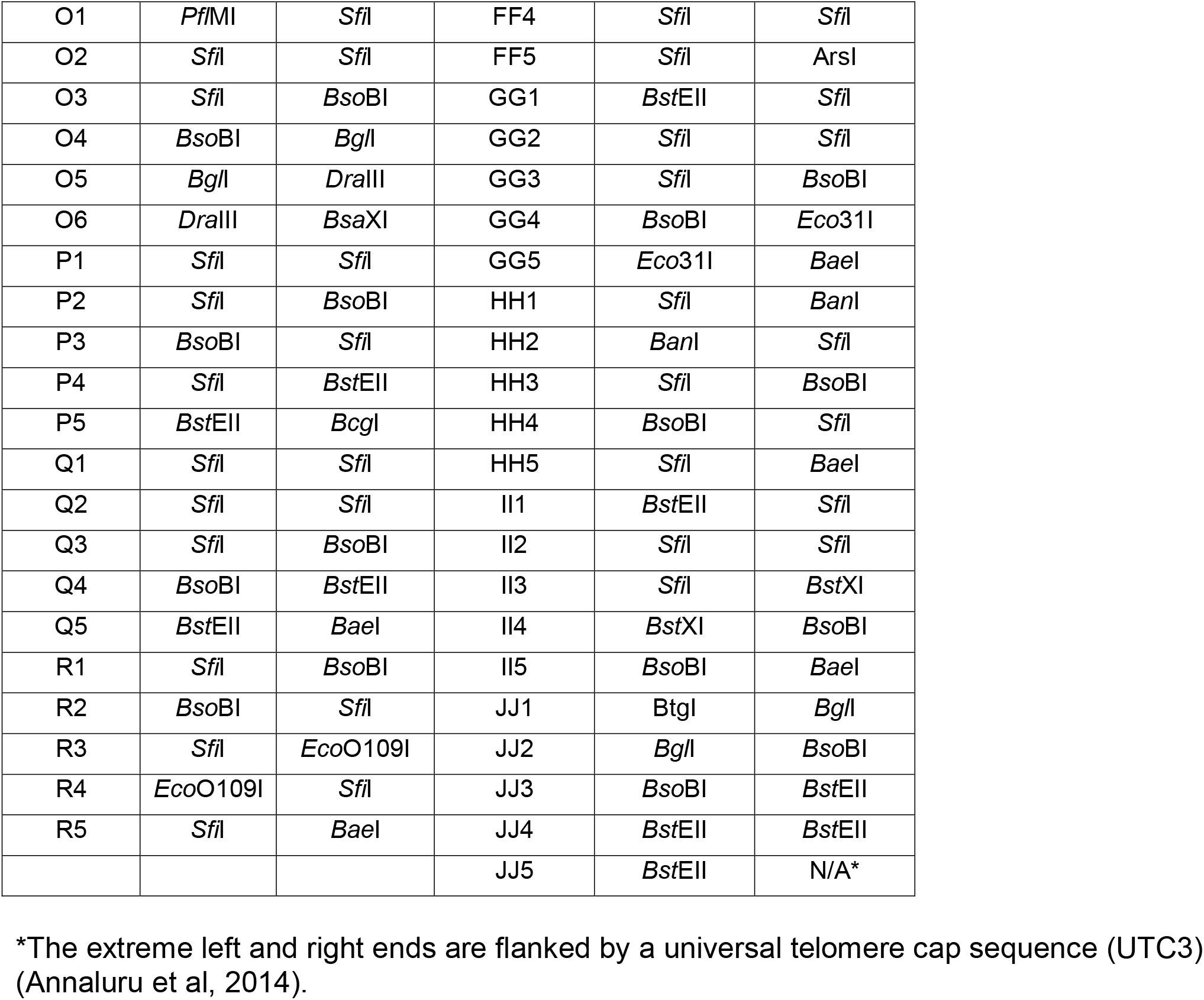
Flanking restriction enzyme sites for all chunks. Related to Figure 1B.

**Table S3.** Flanking restriction enzyme sites for all megachunk clones. Related to Figure 1B.

| Megachunk | Left RE site | Right RE site | Megachunk | Left RE site | Right RE site |
| --- | --- | --- | --- | --- | --- |
| A | *NotI* | *NotI* | S | *FseI* | *FseI* |
| B | *FseI* | *FseI* | T | *NotI* | *NotI* |
| C | *NotI* | *NotI* | U | *NotI* | *NotI* |
| D | *FseI* | *FseI* | V | *NotI* | *NotI* |
| E | *NotI* | *NotI* | W | *FseI* | *FseI* |
| F | *NotI* | *NotI* | X | *NotI* | *NotI* |
| G | *NotI* | *NotI* | Y | *NotI* | *NotI* |
| H | *NotI* | *NotI* | Z | *NotI* | *NotI* |
| I | *NotI* | *NotI* | AA | *NotI* | *NotI* |
| J | *NotI* | *NotI* | BB | *NotI* | *NotI* |
| K | *NotI* | *NotI* | CC | *FseI* | *FseI* |
| L | *NotI* | *NotI* | DD | *NotI* | *NotI* |
| M | *NotI* | *NotI* | EE | *NotI* | *NotI* |
| N | *NotI* | *NotI* | FF | *NotI* | *NotI* |
| O | *NotI* | *NotI* | GG | *NotI* | *NotI* |
| P | *NotI* | *NotI* | HH | *NotI* | *NotI* |
| Q | *NotI* | *NotI* | II | *NotI* | *NotI* |
| R | *NotI* | *NotI* | JJ | *NotI* | *NotI* |

**Table S4.** Strain version table. Related to Figure 3.

| Version name | Strain number | Comment | Details |
| --- | --- | --- | --- |
| yeast_chr04_3_67 | NA | Original design sequence | Final design by BioStudio |
| yeast_chr04_3_68 | NA | Updated design sequence | Modify the repeat smashed essential gene *NOP1* to wild-type *NOP1* with synPCRTags incorporated; shorten the repeat smashed region in *PCF11*. |
| yeast_chr04_9_01 | yWZ572 | synIV Draft strain, with 12 TAG stop codons, 12 loxPsym sites missing, nine wild-type PCRTags (four wild-type PCRTags are the consequence of debugging), 12 point mutations causing amino acid changes, ENA5, ENA2, ENA1 locus duplication. | Remaining TAG stop codons: 220720 T->C, 291609 A->G, 320297 A->G, 334749 A->G, 383360 A->G, 414063 T->C, 439083 T->C, 555123 A->G, 701256 T->C, 954299 T->C, 1015791 T->C, 1148402 T->C. Missing loxPsym sites: 50970-51003, 126870-126903, 169344-169376, 212062-212095, 316483-316516, 334753-334786, 595184-595217, 774040-774073, 776884-776917, 1138870-1138903, 1148365-1148398. Wild-type PCRTags: 401825-401853 YDL019C_2_synR, 587396-587423 YDR083W_1_synF, 593252-593279 YDR087C_1_synR, 671034-671061 YDR126W_1_synF, 775122-775149 YDR172W_2_synF, 777437-777464 YDR173C_1_synF, 811166-811193 YDR192C_1_synR, 874340-874367 YDR227W_1_synF, 981396-981420 YDR289C_1_synR. Point mutations that cause amino acid changes: 6375 G->A, 17617 G->A, 38981 C->A, 80530 A->C, 180759 A->G, 269540 C->T, 695528 G->A, 829793 A->T, 929704 G->A, 979281 G->A, 1330179 G->A, 1385096 C->T. |
| yeast_chr04_9_02 | yWZ608 | Swapped 10 TAG stop codons to TAA in yeast_chr04_9.01, ENA5, ENA2, ENA1 locus duplication. | Remaining TAG stop codons: 383360 A->G, 1148402 T->C. Missing loxPsym sites: 50970-51003, 126870-126903, 169344-169376, 212062-212095, 316483-316516, 334753-334786, 595184-595217, 774040-774073, 776884-776917, 1138870-1138903, 1148365-1148398. Wild-type PCRTags: 401825-401853 YDL019C_2_synR, 587396-587423 YDR083W_1_synF, 593252-593279 YDR087C_1_synR, 775122-775149 YDR172W_2_synF, 777437-777464 YDR173C_1_synF, 811166-811193 YDR192C_1_synR, 874340-874367 YDR227W_1_synF, 981396-981420 YDR289C_1_synR. Point mutations that cause amino acid changes: 6375 G->A, 17617 G->A, 38981 C->A, 80530 A->C, 180759 A->G, 269540 C->T, 695528 G->A, 829793 A->T, 929704 G->A, 979281 G->A, 1330179 G->A, 1385096 C->T. |
| yeast_chr04_9_03 | yWZ628 | Cleared the ENA5, ENA2, ENA1 locus duplication in yeast_chr04_9.02. | Remaining TAG stop codons: 383360 A->G, 1148402 T->C. Missing loxPsym sites: 50970-51003, 126870-126903, 169344-169376, 212062-212095, 316483-316516, 334753-334786, 595184-595217, 774040-774073, 776884-776917, 1138870-1138903, 1148365-1148398. Wild-type PCRTags: 401825-401853 YDL019C_2_synR, 587396-587423 YDR083W_1_synF, 593252-593279 YDR087C_1_synR, 775122-775149 YDR172W_2_synF, 777437-777464 YDR173C_1_synF, 811166-811193 YDR192C_1_synR, 874340-874367 YDR227W_1_synF, 981396-981420 YDR289C_1_synR. Point mutations that cause amino acid changes: 6375 G->A, 17617 G->A, 38981 C->A, 80530 A->C, 180759 A->G, 269540 C->T, 695528 G->A, 829793 A->T, 929704 G->A, 979281 G->A, 1330179 G->A, 1385096 C->T. |
| yeast_chr04_9_04 | yWZ647 | Swapped 775122-775149 YDR172W_2_synF and 777437-777464 | Remaining TAG stop codons: 383360 A->G, 1148402 T->C. Missing loxPsym sites: 50970-51003, 126870-126903, 169344-169376, 212062-212095, 316483-316516, 334753-334786, 595184-595217, 774040-774073, 776884-776917, 1138870-1138903, 1148365-1148398. Wild- |
|  |  | YDR173C_1_synF<br>PCRTags to synthetic in<br>yeast_chr04_9_04. | type PCRTags: 401825-401853 YDL019C_2_synR, 587396-587423 YDR083W_1_synF, 593252-593279 YDR087C_1_synR, 811166-811193 YDR192C_1_synR, 874340-874367 YDR227W_1_synF, 981396-981420 YDR289C_1_synR. Point mutations that cause amino acid changes: 6375 G->A, 17617 G->A, 38981 C->A, 80530 A->C, 180759 A->G, 269540 C->T, 695528 G->A, 829793 A->T, 929704 G->A, 979281 G->A, 1330179 G->A, 1385096 C->T. |
| yeast_chr04_9_05 | yWZ674 | Swapped two more TAG<br>stop codons to TAA in<br>yeast_chr04_9.03.<br>Inserted two loxPsym sites<br>that are missing in<br>yeast_chr04_9.03.<br>Introduced a 2078 bp<br>deletion in megachunk F<br>by two loxPsym sites<br>recombination | Missing loxPsym sites: 50970-51003, 126870-126903, 169344-169376, 212062-212095, 226643-226676, 316483-316516, 595184-595217, 774040-774073, 1138870-1138903, 1148365-1148398. Wild-type PCRTags: 401825-401853 YDL019C_2_synR, 587396-587423 YDR083W_1_synF, 593252-593279 YDR087C_1_synR, 775122-775149 YDR172W_2_synF, 777437-777464 YDR173C_1_synF, 811166-811193 YDR192C_1_synR, 874340-874367 YDR227W_1_synF, 981396-981420 YDR289C_1_synR. Point mutations that cause amino acid changes: 6375 G->A, 17617 G->A, 38981 C->A, 80530 A->C, 180759 A->G, 269540 C->T, 695528 G->A, 829793 A->T, 929704 G->A, 979281 G->A, 1330179 G->A, 1385096 C->T. Deletion region between two loxPsym sites: 220720-222798. |
| yeast_chr04_9_06 | yWZ687 | Fixed the 2078 bp deletion<br>in megachunk F of<br>yeast_chr04_9.04. | Missing loxPsym sites: 50970-51003, 126870-126903, 169344-169376, 212062-212095, 226643-226676, 316483-316516, 774040-774073, 1138870-1138903. Wild-type PCRTags: 401825-401853 YDL019C_2_synR, 587396-587423 YDR083W_1_synF, 593252-593279 YDR087C_1_synR, 775122-775149 YDR172W_2_synF, 777437-777464 YDR173C_1_synF, 811166-811193 YDR192C_1_synR, 874340-874367 YDR227W_1_synF, 981396-981420 YDR289C_1_synR. Point mutations that cause amino acid changes: 6375 G->A, 17617 G->A, 38981 C->A, 80530 A->C, 180759 A->G, 269540 C->T, 695528 G->A, 829793 A->T, 929704 G->A, 979281 G->A, 1330179 G->A, 1385096 C->T. |
| yeast_chr04_9_07 | yWZ703 | Fixed the 38981 C->A<br>mutation in<br>yeast_chr04_9_06. | Missing loxPsym sites: 50970-51003, 126870-126903, 169344-169376, 212062-212095, 226643-226676, 316483-316516, 774040-774073, 1138870-1138903. Wild-type PCRTags: 401825-401853 YDL019C_2_synR, 587396-587423 YDR083W_1_synF, 593252-593279 YDR087C_1_synR, 775122-775149 YDR172W_2_synF, 777437-777464 YDR173C_1_synF, 811166-811193 YDR192C_1_synR, 874340-874367 YDR227W_1_synF, 981396-981420 YDR289C_1_synR. Point mutations that cause amino acid changes: 6375 G->A, 17617 G->A, 80530 A->C, 180759 A->G, 269540 C->T, 695528 G->A, 829793 A->T, 929704 G->A, 979281 G->A, 1330179 G->A, 1385096 C->T. |

**Table S5.**
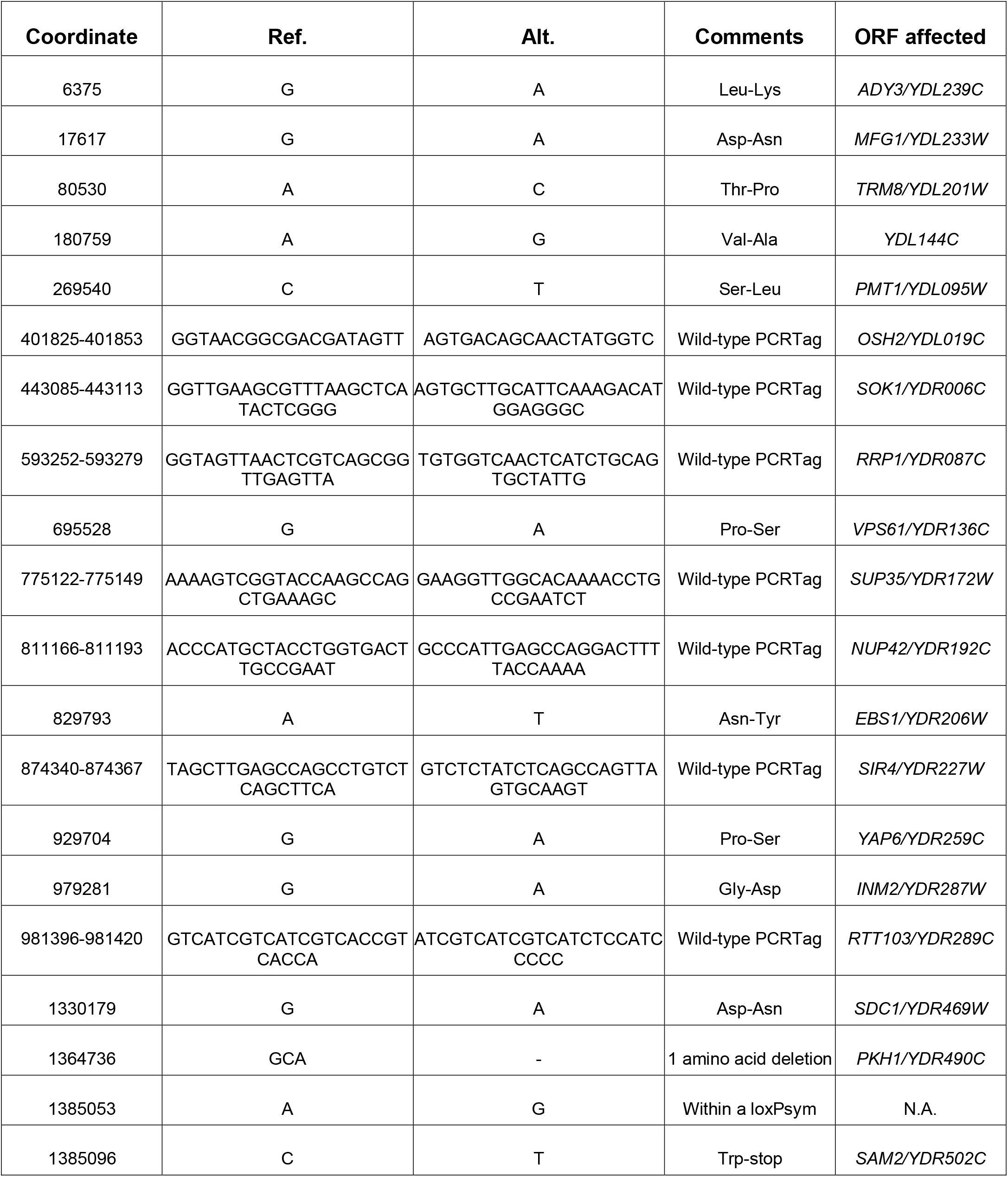
SNPs and indels in yeast_chr04_9_07 strain. Related to Figure 3.

**Table S6.**
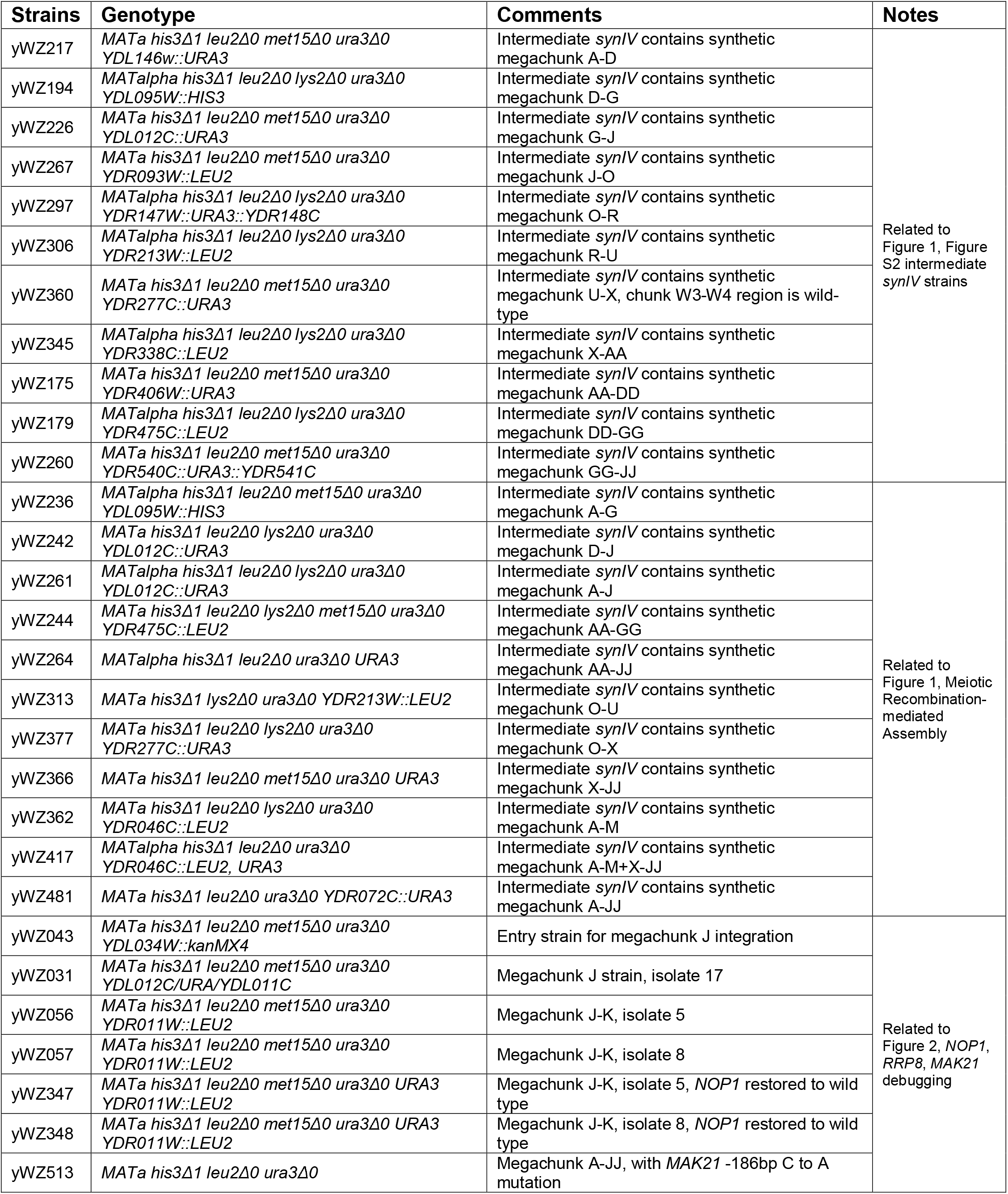
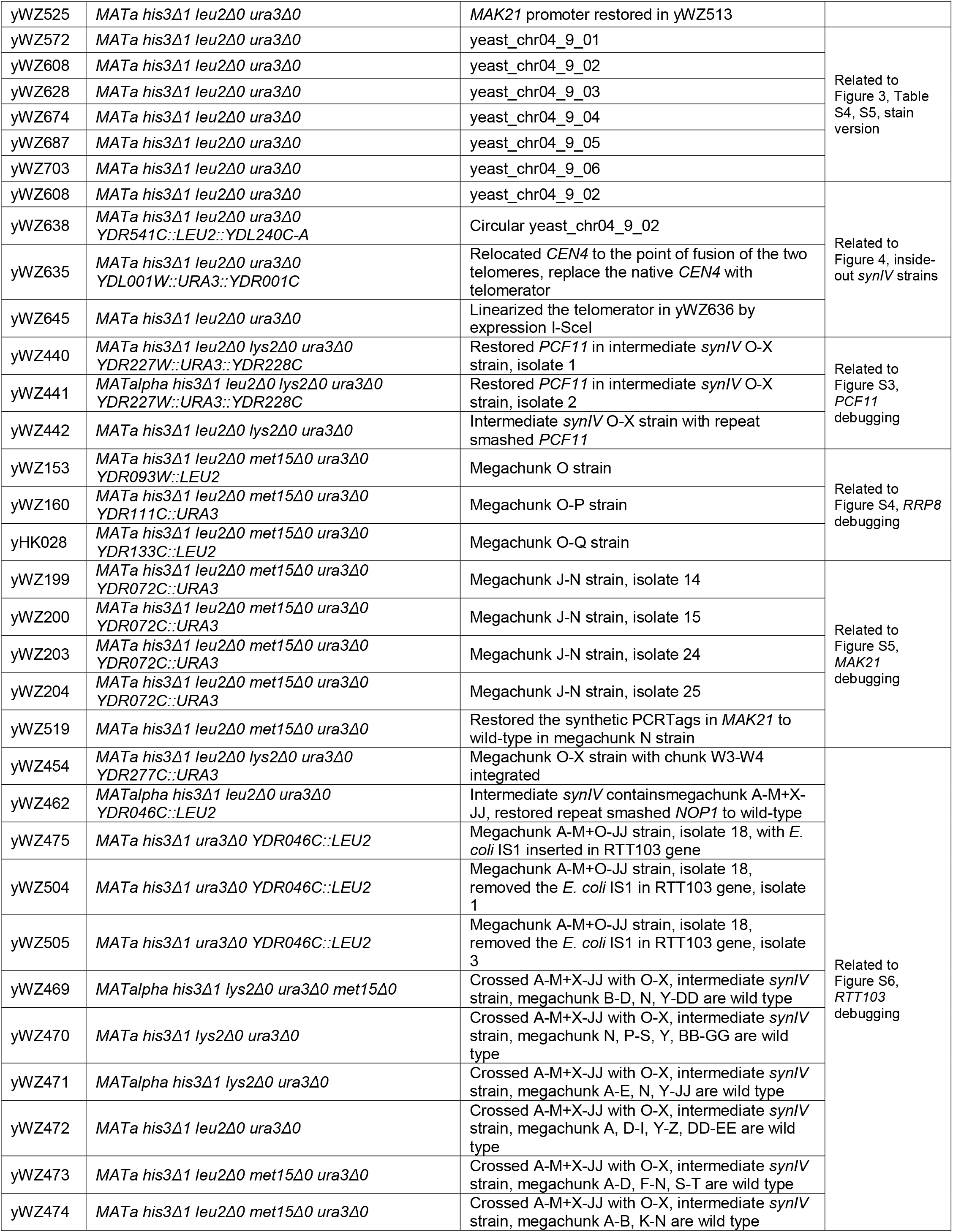
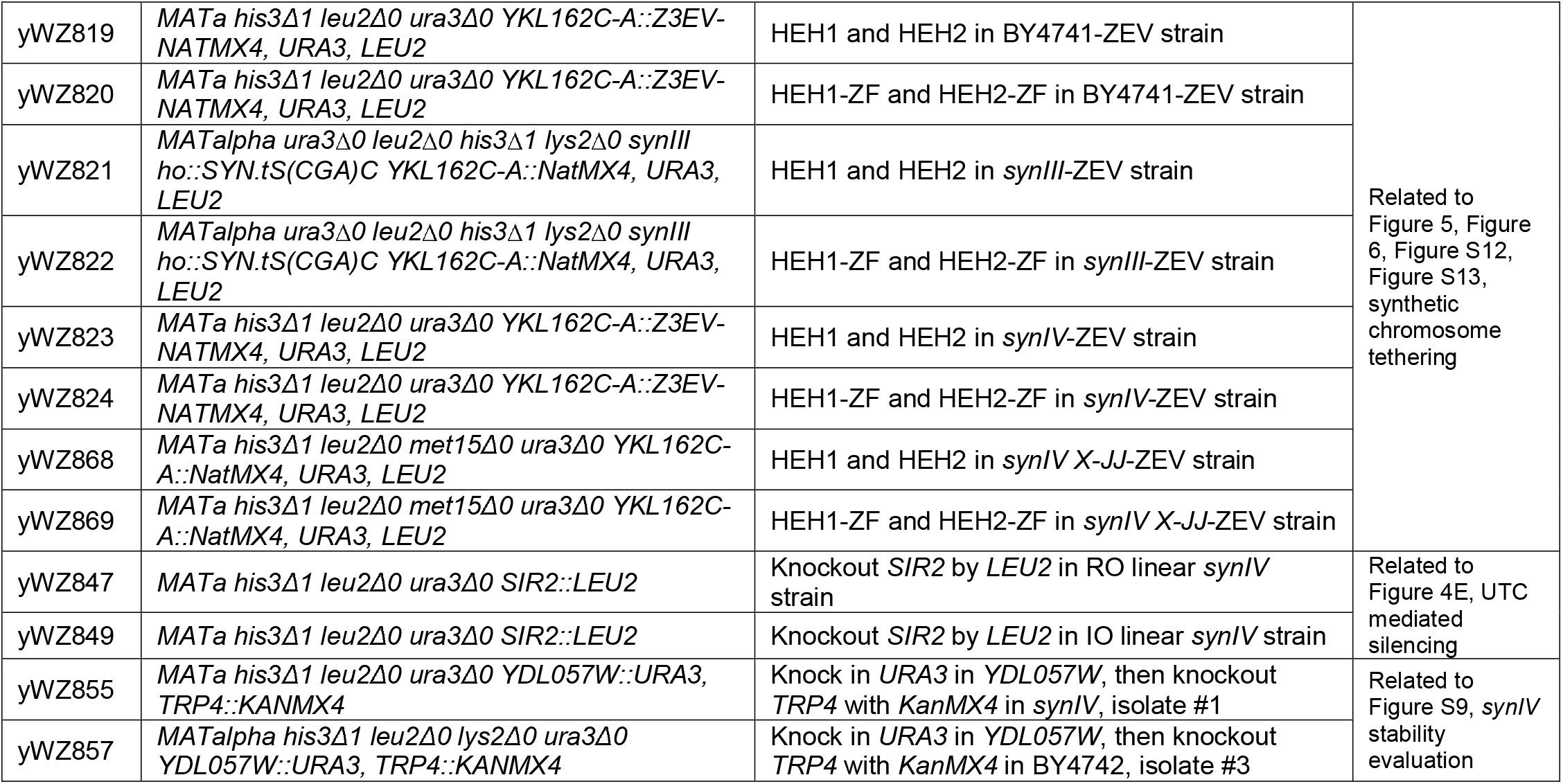
Strains used in this study.

